# Aneuploid cells activate NF-κB to promote their immune clearance by NK cells

**DOI:** 10.1101/2020.06.25.172239

**Authors:** Ruoxi W. Wang, Sonia Viganò, Uri Ben-David, Angelika Amon, Stefano Santaguida

**Author notes:** Correspondence to (SS).

## Abstract

The immune system plays a major role in the protection against cancer. Identifying and characterizing the pathways mediating this immune surveillance is thus critical for understanding how cancer cells are recognized and eliminated. We previously found that untransformed cells that had undergone senescence due to highly abnormal karyotypes are eliminated by Natural Killer (NK) cells *in vitro*. Here we show that this is also true for aneuploid untransformed cells that had not lost their ability to proliferate. Their elimination by NK cells, like that of aneuploid senescent cells, is predominantly mediated by non-cell autonomous mechanisms. Our data further indicate that NF-κB signaling in aneuploid cells is central to eliciting this immune response. Inactivating NF-κB abolishes NK-cell mediated clearance in aneuploid cells *in vitro*. In cancer cell lines, NF-κB signaling correlates with degree of aneuploidy, raising the possibility that aneuploidy-induced immune recognition is partially retained in cancer.

## Introduction

Aneuploidy is defined as a state in which the chromosome number is not a multiple of the haploid complement (Pfau & Amon 2012). In all organisms analyzed to date, an unbalanced karyotype has detrimental effects (Pfau & Amon 2012; Santaguida & Amon 2015). In yeast, aneuploidy leads to proliferative defects and proteotoxic stress (Torres et al. 2010). The impact of aneuploidy on higher eukaryotes is even more severe. Most single autosomal gains and all autosomal losses cause embryonic lethality. The aneuploidies that do survive embryonic development cause significant anatomical and physiological abnormalities. This severe impact of aneuploidy on mammalian physiology is also reflected at the cellular level. Trisomic mouse embryonic fibroblasts (MEFs) and aneuploid human cells proliferate more slowly compared to their euploid counterparts and experience a variety of cellular stresses (Williams et al. 2008; Pfau et al. 2016; Santaguida et al. 2015). One of the stresses, elicited by aneuploidy, is replication stress. Aneuploidy-induced replication stress triggers genome instability, thereby fueling the evolution of highly abnormal karyotypes (Ohashi et al. 2015; Lamm et al. 2016; Sheltzer et al. 2012; Santaguida et al. 2017). Recently, we showed that cells harboring such aberrant karyotypes are recognized and eliminated by Natural Killer (NK) cells *in vitro* (Santaguida et al. 2017).

Although aneuploidy is highly detrimental at both the cellular and organismal level in untransformed cells, it is also a hallmark of cancer, a disease characterized by uncontrolled cell proliferation (Gordon et al. 2012). About 90% of solid tumors and 75% of hematopoietic malignancies are characterized by whole chromosome gains and losses (Weaver & Cleveland 2006). A high degree of aneuploidy is often associated with poor prognosis, immune evasion and a reduced response to immunotherapy (Ben-David & Amon 2019). Given the negative effects of aneuploidy on primary cells, it remains unclear how cells with severe genomic imbalances gain tumorigenic potential. Furthermore, which aneuploidy-associated molecular features alter immune recognition during tumor evolution remains an active field of research.

By inducing high levels of chromosome mis-segregation followed by continuous culturing, cells with abnormal complex karyotypes are generated that eventually cease to divide and enter a senescent-like state. We have named such cells <u>Ar</u>rested with <u>C</u>omplex <u>K</u>aryotypes (ArCK) cells (Santaguida et al. 2017; Wang et al. 2018). Previous work indicated that ArCK cells upregulate gene expression signatures related to an immune response that render them susceptible to elimination by Natural Killer (NK) cells *in vitro* (Santaguida et al. 2017). However, the molecular and functional bases for this immune recognition of ArCK cells remained unclear. Several pathways could be involved in this process. Nuclear factor-kappaB (NF-κB) is induced under several stress conditions to elicit a pro-inflammatory response (Hayden & Ghosh 2012; T. Liu et al. 2017). In the canonical NF-κB pathway, upon stress induction, IκB kinase complex (IKK) phosphorylates IκB, thereby marking it for proteolytic degradation (Perkins 2007). As a result of this degradation, RelA-p50 translocates into the nucleus where it activates expression of pro-inflammatory genes. In the non-canonical NF-κB pathway, phosphorylation and cleavage of p100 triggers the nuclear translocation of the RelB-p52 complex to induce a pro-inflammatory response (Zhang et al. 2017). Recent studies further suggest that cytosolic nucleic acids lead to cGAS/STING activation in senescent cells that induces an interferon response via JAK-STAT signaling pathway (Dou et al. 2017; Vizioli et al. 2020).

Here we investigate which innate immune pathways contribute to NK cell-mediated elimination of aneuploid cells. We show that both the canonical and non-canonical NF-κB pathways generate pro-inflammatory signals in ArCK cells and aneuploid cells that have retained their ability to proliferate. Inactivating both NF-κB pathways in cells harboring an unbalanced karyotype prevents NK cell-mediated killing *in vitro*. Consistent with aberrant karyotypes eliciting an NF-κB response, we further find that this pro-inflammatory gene expression signature is present in highly aneuploid human cancer cell lines. Our data are the first to provide insight into the pathways mediating an aneuploidy-induced immune response.

## Results

### An assay to assess elimination of ArCK cells by natural killer (NK) cells *in vitro*

To address the molecular basis for immune recognition of ArCK cells, we established a coculture system to monitor the interactions between NK cells and ArCK cells. In this setup, we utilized human, untransformed RPE1-hTERT cells in which chromosome segregation errors were induced by inhibiting the function of the spindle assembly checkpoint (SAC) (Santaguida et al. 2010). We synchronized cells in G1 then released them into the cell cycle in the presence of the SAC kinase inhibitor reversine (Figure S1A). We removed the drug after cells had undergone an aberrant mitosis due to SAC inhibition. 72 hours after inducing chromosome mis-segregation, we exposed cells for 12 hours to the spindle poison nocodazole. This arrested dividing cells in mitosis and allowed us to remove them from cell culture by mitotic shake-off (Wang et al. 2018). To remove all proliferating cells, we repeated this nocodazole treatment and shake-off procedure 4 more times. Cells that remained on the tissue culture plate by the end of the procedure harbored highly aberrant karyotypes and had entered a senescent-like state (Figure S1A; (Santaguida et al. 2017)). We then co-cultured these ArCK cells with an immortalized NK cell line activated by constitutive IL2 expression (NK92-MI; (Tam et al. 1999)) and monitored their interactions by live cell imaging.

To be able to quantify the degree of NK cell killing of ArCK cells, we established the following criteria. We defined a killing event when an ArCK cell was (1) engaged by one or multiple NK cells and (2) was lifted from the tissue culture plate (Figure 1A). We chose these criteria because they coincided with membrane permeabilization as judged by the ability of the nucleic acid dye TO-PRO3 to enter a cell (Figure 1A). We tracked individual target cells and recorded the time when each of them was killed during the 36-hour live cell imaging. If a cell divided during the time course, we followed only one of the resulting two cells for the remainder of the assay. We then calculated cumulative cell death for each condition and generated killing curves at hourly resolution. We found that at a ratio of 2.5 NK cells to 1 ArCK cell, ArCK cells were consistently killed two times more effectively during a 36-hour co-culture experiment than euploid control cells (Figure 1B).

**Figure 1.**
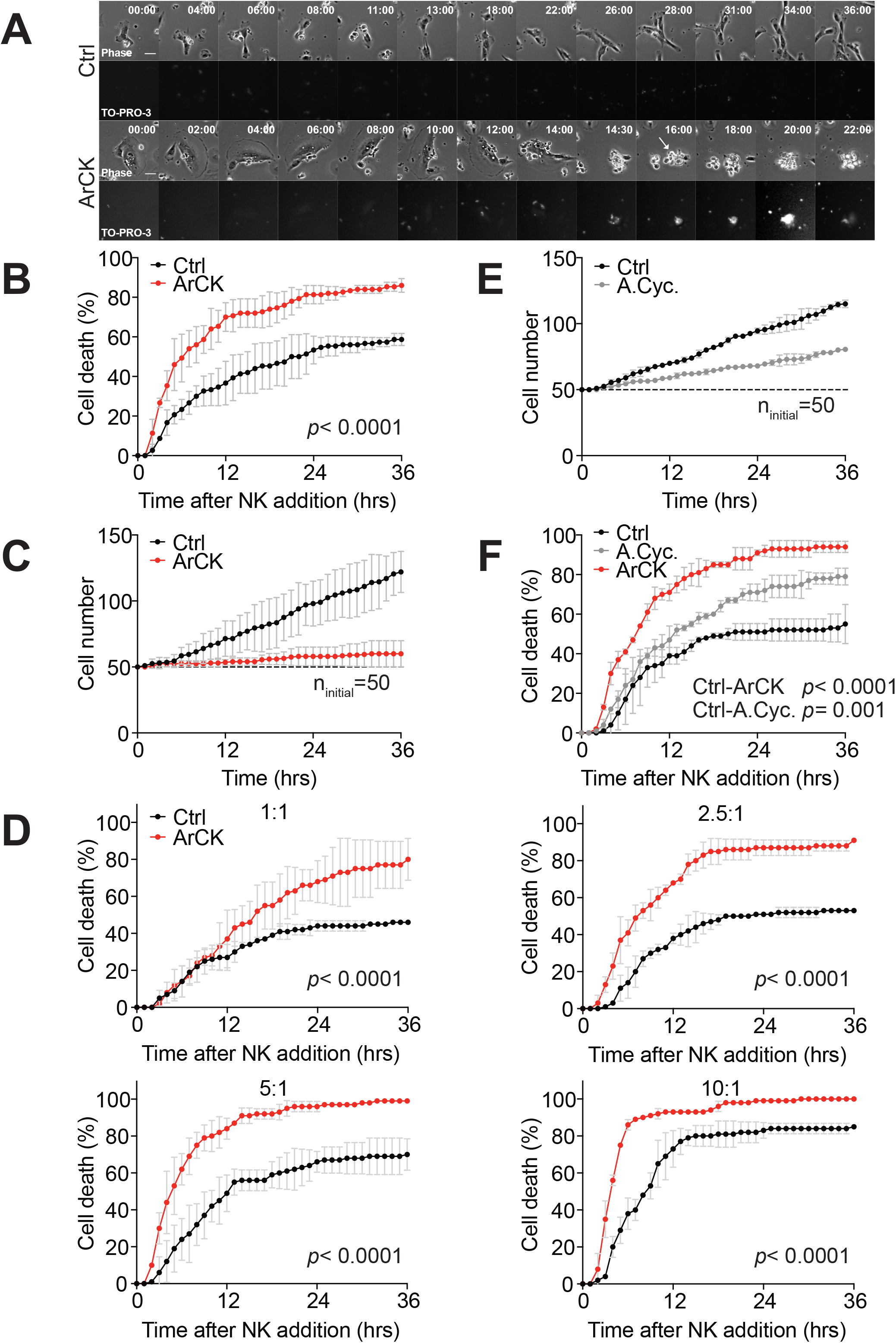
Aneuploid cells are recognized by Natural Killer (NK) cells *in vitro*. (A) Representative images of euploid and ArCK cells interacting with NK cells. To generate ArCK cells, RPE1-hTERT cells were synchronized at the G1/S border by thymidine block for 24 hours. 6 hours after thymidine release, cells were treated with reversine (500 nM) for 18 hours. 60 hours after reversine washout, which corresponds to roughly 72 hours after inducing chromosome mis-segregation, cells were treated with nocodazole (100 ng/mL) for 12 hours. Dividing cells were then removed from the plate by mitotic shake-off. A total of five shake-offs were performed at a 12-hour interval to ensure complete removal of proliferating cells from the population. The cells left on the plate after five shake-offs were collected and designated as the ArCK sample. To measure NK cell-mediated cytotoxicity, target cells were pre-incubated in NK cell medium for 10 hours before adding NK cells at a 2.5:1 (NK cells: target cells) ratio. NK cell-mediated killing was recorded by live cell imaging for 36 hours at a 30-min interval. TO-PRO-3 (1 μM) was added to the medium at the same time of NK addition to measure cell membrane integrity. Phase contrast (top) and TO-PRO-3 signal (bottom) from the same field are shown. The arrow indicates ArCK cell death. All images were acquired at the same exposure time and light intensity. Scale bar 20 μM. (B) Measurement of NK cell-mediated killing of ArCK and euploid control cells (Ctrl). 50 randomly chosen cells were grown as described in (A) and followed for 36-hours by live cell imaging. A cell is defined as being killed when the target cell is fully engaged by a single or multiple NK cell(s) and completely lifted from the culture plate. When a cell divided during the analysis, only one of the two cells was followed after cell division. Cumulative cell death was calculated and plotted at an hourly resolution (n=3; mean ± SD). To determine statistical significance, the times when a target cell was killed by NK cells were pooled in each condition and the significance was determined by nonparametric Kolmogorov-Smirnov test (KS test). (C) Measurement of ArCK cell proliferation without NK cells. Live cell imaging of target cells without NK cells was performed using the same condition as described in (A). 50 cells were randomly chosen at the beginning of the movie as the initial population (indicated by the dashline, n_initiai_ = 50). The cumulative cell number was recorded and plotted at an hourly resolution for 36 hours (n=2; mean ± SD). (D) NK cell-mediated cytotoxicity across various NK cell to target cell ratios. Either normal euploid (Ctrl) or ArCK cells were co-cultured with NK cells at the indicated NK cell: target cell ratios. The cumulative killing of target cells is measured as described in (B; n=2; mean ± SD). Statistical analyses were performed as in (B). (E) Cell proliferation in aneuploid cycling cells. RPE1-hTERT cells were synchronized at the G1/S border by thymidine block for 24 hours. 6 hours after release from the thymidine block, cells were treated with reversine (500 nM) for 18 hours. 48 hours after reversine washout, corresponding to about 60 hours after the faulty mitosis, cells were collected and were referred to as aneuploid cycling cells (A.Cyc.). Cell proliferation during the 36-hour time lapse was measured as described in (C). The dashed line indicates the starting cell number (ninitial=50; n=2; mean ± SD). (F) NK cell-mediated killing of euploid proliferating (Ctrl), aneuploid cycling (A. Cyc.), and ArCK cells. Cells were grown and analyzed within the same experiment (n=3; mean ± SD). Statistical analyses were performed as in (B).

ArCK cells are largely senescent (Santaguida et al. 2017). Hence, they hardly divided during the 36-hour time lapse employed in our NK cell killing assay (Figure 1C). In contrast, euploid cells continued to divide. It was thus possible that the difference in NK cell-mediated cytotoxicity towards euploid and ArCK cells was because NK cells became limiting when co-cultured with euploid cells but not with aneuploid cells. To test this possibility, we analyzed the effects of changing the ratio of NK cells and ArCK cells on NK cell-mediated killing. We found that even at high NK cell to ArCK cell ratio (5:1 and 10:1), aneuploid cells were still more effectively killed than euploid cells (Figure 1D). We conclude that NK cells are not limiting in our assay. We further note that when a cell divided during observation, we followed only one of the two cells after cell division, which corrects for the bias in target cell number. To address the possibility that NK cells become exhausted during the course of the co-culture experiment, we divided the 36-hour assay into two time courses, where the same population of NK cells was consecutively co-cultured with ArCK cells for 18 hours each. NK cells were equally effective in killing ArCK cells (Figure S1B) in this experimental setup, indicating that NK cell exhaustion did not occur within the course of the analysis. We propose that the eventual plateauing of killing as the assay proceeds is likely due to NK cells taking longer to find their targets.

We next set out to test why euploid cells are readily killed by NK cells in our *in vitro* assay. One possible explanation was that RPE1-hTERT cells express human telomerase reverse transcriptase (hTERT), which may generate oncogenic transformation associated NK cell stimulatory signals (Chiossone et al. 2018; Shimasaki et al. 2020). To test whether this was the case, we assessed NK cell-mediated cytotoxicity across three different types of early passage euploid primary fibroblasts (human embryonic lung fibroblast, IMR90; normal neonatal or adult human dermal fibroblasts, NHDF-Neo or NHDF-Ad). DNA sequencing indicated that both neonatal or adult human dermal fibroblasts remained stably euploid for at least 15 passages (data not shown). The analysis of these primary cells revealed large variations in NK cell-mediated killing (Figure S1C and D). Adult human dermal fibroblasts were not readily eliminated by NK-cells whereas both neonatal human dermal fibroblasts and IMR90 cells were highly immunogenic. It thus appears that NK cell-mediated killing differs significantly between primary cultured cells. Given this variability, we decided to focus on investigating the effects of karyotype alterations on NK-cell interactions using RPE1-hTERT cells, as we had already developed protocols to generate aneuploid and euploid cell populations using this cell type. We conclude that in the assay we developed here, highly aneuploid senescent RPE1-hTERT cells are more effectively recognized and eliminated by NK92-MI cells *in vitro* than their euploid counterparts.

### Aneuploid cycling cells are recognized and eliminated by NK cells *in vitro*

Previous studies suggested that immediately after reversine-induced chromosome missegregation, ~90% of cells harbor single chromosome gains or losses. This degree of aneuploidy does not preclude them from proliferating, but they do so more slowly ((Santaguida et al. 2017; Santaguida et al. 2015); Figure 1E, S1E). To determine whether aneuploid cells that have not yet lost their ability to divide are also recognized and eliminated by NK cells *in vitro*, we induced chromosome mis-segregation using reversine and collected cells ~60 hours thereafter. We will refer to these cells as “aneuploid cycling cells”. Single cell sequencing analysis revealed that more than 40% of the cells treated in this manner harbor random chromosome gains and/or losses (Figure S1F and G). Aneuploid cycling cells were recognized and eliminated by NK cells, although to a lesser degree than ArCK cells (Figure 1F). We conclude that cellular senescence is not a prerequisite for aneuploid cell recognition by NK cells *in vitro*.

### Prolonged G1 arrest associated with cell size increase elicits NK cell-mediated cytotoxicity

Our results showed that ArCK cells were more effectively killed by NK cells compared to aneuploid cycling cells. We thus hypothesized that G1 arrest contributes to NK cell-mediated killing. Indeed, permanent cell-cycle arrest has been shown to elicit an immune response (Gorgoulis et al. 2019). To determine whether G1 arrest *per se* is sufficient to cause immune recognition, we assessed NK cell-mediated cytotoxicity towards G1 arrested cells induced by three different methods. We treated RPE1-hTERT cells for 7 days with 1) the topoisomerase II inhibitor, doxorubicin, to induce high levels of DNA double-strand breaks (Pommier et al. 2010); 2) the cyclin-dependent kinases CDK4/6 inhibitor, palbociclib; or 3) the imidazoline analog, nutlin3, to disrupt the interaction between p53 and its ubiquitin ligase Mdm2 thereby stabilizing p53.

DNA content analysis by flow cytometry showed that after 7 days of treatment, cells were arrested in G1 (Figure 2A; note that the G1 peaks are unusually broad. This is because cells continue to grow in size during the G1 arrest (Figure 2B), which leads to an increase in mitochondrial DNA, causing the G1 peak to broaden (Levy & Heald 2012; Neurohr et al. 2018)). With the exception of nutlin3 treatment, these G1 arrests were irreversible: cells did not resume proliferation following drug washout as judged by EdU incorporation and cell proliferation assays (Figure 2C and D). Co-culturing these large G1-arrested cells with NK cells revealed that irrespective of the means by which the arrest was induced, NK cells exhibit ~2 fold higher killing on these large G1 arrested cells compared to the untreated proliferating control cells (Figure 2E).

**Figure 2.**
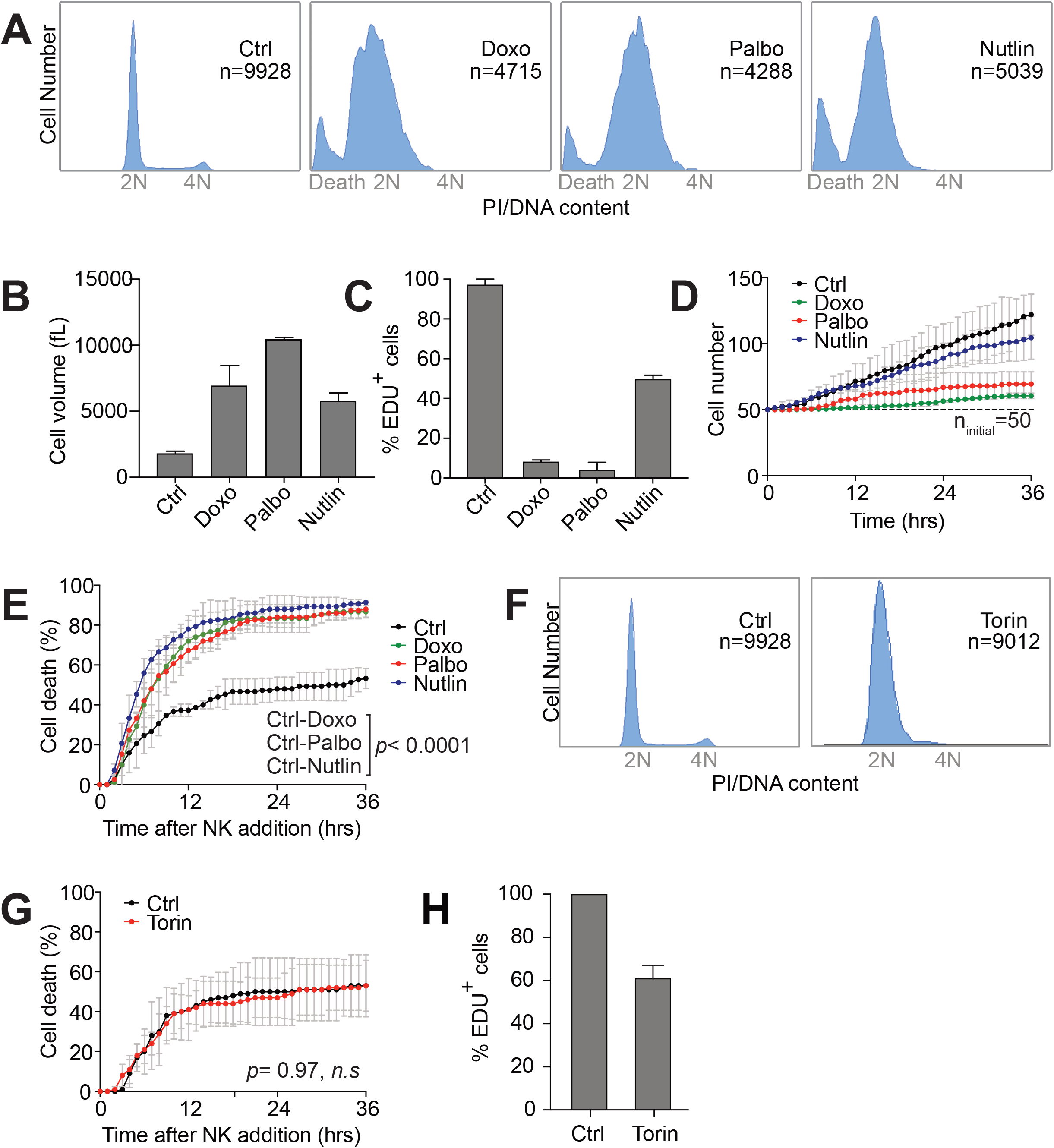
Prolonged G1 arrest associated with cell size increase elicits NK cell-mediated cytotoxicity. (A) DNA content analysis of various G1 arrests. RPE1-hTERT cells were treated for 7 days with doxorubicin (Doxo; 100 ng/ml), palbociclib (Palbo; 5 μM), or nutlin3 (Nutlin; 10 μM). Total number of cells analyzed is indicated by *n* in each condition (n=2). Results were comparable between replicates. (B) Cell size analysis for doxorubicin (Doxo), palbociclib (Palbo), or nutlin 3 (Nutlin) treated cells. The population local mode in each condition was determined (n ≥□5000 for each condition; mean ± SD). (C) EdU incorporation following drug washout. RPE1-hTERT cells were treated with doxorubicin (Doxo), palbociclib (Palbo), or nutlin 3 (Nutlin) for 7 days as described in (A). Cells were then washed three times with PBS and incubated with medium containing 5-ethynyl-2’-deoxyuridine (EdU; 10 μM) for 48 hours. The percentage of EdU positive cells was determined (n ≥□100 for each condition; mean ± SD). (D) Cell proliferation following release from various G1 arrests. After 7 days of doxorubicin (Doxo), palbociclib (Palbo), or nutlin3 (Nutlin) treatment, the drugs were washed out and cells were re-plated for live cell imaging. Cell proliferation was measured as described in Figure 1D. The dashed line indicates the starting cell number (n_initial_ = 50; n=2; mean ± SD). (E) NK cell-mediated killing for doxorubicin (Doxo), palbociclib (Palbo), and nutlin 3 (Nutlin) treated samples (NK cell: target cell =2.5:1; n=3; mean ± SD). Statistical analyses were performed as in Figure 1B. (F) DNA content analysis of torin1 arrested cells. RPE1-hTERT cells were treated torin 1 (Torin; 5 μM) for 24 hours. Total number of cells analyzed is indicated by *n* in each condition. Because the torin 1 treated sample was analyzed at the same time as samples described in (A), the same control panel (Ctrl) is shown (n=2). Results were comparable between replicates. (G) NK cell-mediated cytotoxicity towards torin1-treated cells. Torin1 treated (Torin) cells were generated as described in (F) and the NK cell killing assay was performed as described in Figure 1A (n=2; mean ± SD). Statistical analyses were performed as in Figure 1B (*n.s* – not significant). (H) Cell proliferation after torin1 washout. Cells were treated with torin1 (5 μM) for 24 hours and released into medium lacking the drug but containing EdU (10 μM) for 48 hours. The percentage of EdU positive cells was determined (n ≥□100 for each condition; mean ± SD).

Inactivation of the TORC1 pathway also causes cells to arrest in G1 (Sousa-Victor et al. 2015), but cells neither increase in size during the arrest nor undergo senescence (Sousa-Victor et al. 2015; Kucheryavenko et al. 2019). RPE1-hTERT cells were arrested in G1 by treating them with 5 μM of the mTOR kinase inhibitor torin1, for 24 hours (Figure 2F). Yet NK cell recognition and killing was not significantly affected by torin treatment (Figure 2G). This was not due to the fact that torin1-treated cells resumed proliferation when the drug was removed prior to coculturing with NK cells (note drug removal was necessary to ensure that they do not interfere with NK cell function in the co-culture assay). EdU incorporation analysis showed that the degree of cell division, following drug removal was similar between torin1- and nutlin3-treated cells (compare Figures 2C and 2H). We conclude that G1 arrest in target cells contributes to NK cell engagement, but only when accompanied by features of senescence.

### Aneuploidy causes senescence-independent NK cell recognition

The observation that senescence triggered by multiple mechanisms caused NK cell recognition begged the question of whether aneuploid cells are recognized and eliminated simply because they entered a terminal G1 arrest. To address this possibility, we compared the degree of senescence in ArCK and aneuploid cycling cells with that of cells treated with doxorubicin, palbociclib or nutlin3 for 7 days. First, we assessed DNA damage levels across all conditions by measuring nuclear γ-H2AX foci (Figure 3A, S2A). DNA damage will increase the expression of NK cell activating receptor (NKG2D) ligands such as MICA and ULBP2, thereby triggering NK cell-mediated clearance (Raulet et al. 2013). In untreated control proliferating cells, more than 80% of the cells harbor fewer than 10 γ-H2AX foci per nucleus (Figure 3A, S2A). As expected, doxorubicin caused high levels of DNA damage-about 90% of the cells displayed more than 20 foci and amongst them more than 50% of the cells harbored 50 foci or more. In contrast, palbociclib and nutlin3 treated cells did not exhibit significant increase in DNA damage compared to untreated control cells. About one third of ArCK cells harbored more than 10 foci, likely caused by replication stress and/or endogenous reactive oxygen species (ROS) associated with aneuploidy (Santaguida et al. 2017; M. Li et al. 2010). Aneuploid cycling cells did not experience significant DNA damage within the nucleus (Figure 3A). However, these cells harbored micronuclei, the result of chromosomes left behind during the reversine-induced faulty cell division (Figure 3B). As observed previously, micronuclei in aneuploid cycling cells harbor DNA damage as indicated by bright γ-H2AX foci (Figure S2A; (Crasta et al. 2012)).

**Figure 3.**
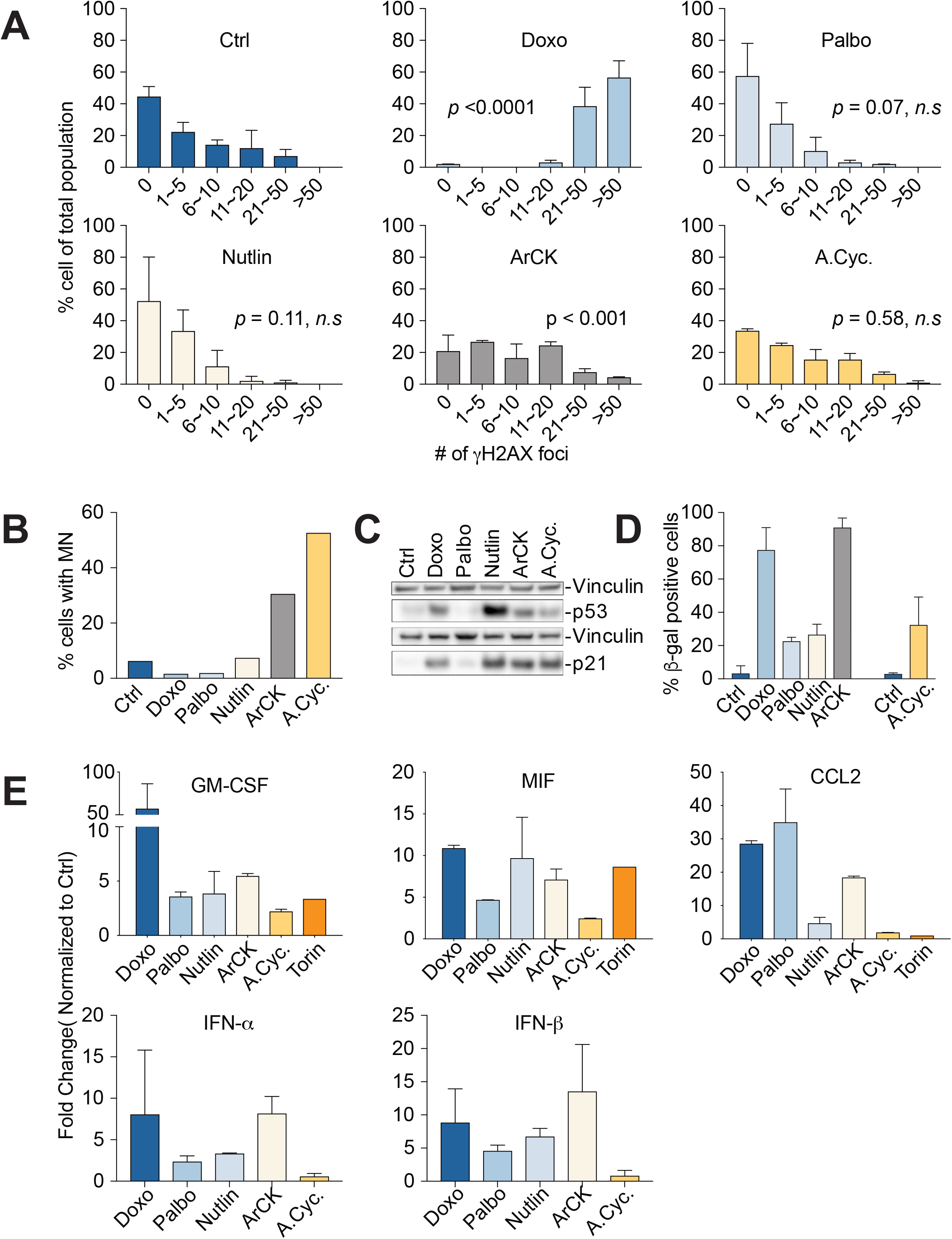
Aneuploidy causes NK cell recognition by mechanisms in addition to triggering senescence. (A) γ-H2AX were analyzed in ArCK, aneuploid cycling (A.Cyc., generated as described in Figure 1) and G1 arrested cells (generated as described in Figure 2). Only foci in the main nucleus were counted (n≥⍰50; mean ± SD). The distribution of focus number in each condition was compared to that of control based on the raw focus number counting using the Kolmogorov-Smirnov test (*n.s* – not significant). (B) The percentage of cells harboring micronuclei was analyzed on the samples generated as described in (A; n ≥100 for each condition). (C) p53 and p21 levels were determined by western blot analysis on the samples generated as described in (A). Vinculin was used as loading control. Results were comparable between replicates. (D) The degree of senescence was measured by senescence-associated β-galactosidase (β-gal) activity on samples generated as described in (A). The graph shows the percentage of β-gal positive cells (n > 100). (E) Secreted cytokine and interferon levels were determined in cell supernatants. Media were collected after 36 hours of incubation with cells grown as described in (A). Cytokine and interferon levels are shown as fold change of euploid control cells (mean ± SD).

Induction of the DNA damage response genes p53 and p21 matched the degree of DNA damage in various cells with the obvious exception of nutlin3 treated cells (Figure 3C). Senescence-associated beta-galactosidase activity (SA-beta-gal), a biomarker frequently used to assess degree of senescence, also agreed with the degree of nuclear DNA damage (Figure 3D, S2B). In doxorubicin treated and ArCK cells, a high degree of beta-gal positive cells was observed. Palbociclib-treated, nutlin3-treated, and aneuploid cycling cells exhibited only slightly increased levels of SA-beta-gal positive cells compared to the untreated proliferating control cells (Figure 3D, S2B).

We then examined the senescence associated secretory phenotype (SASP), which plays a critical role in immune cell recruitment (Gorgoulis et al. 2019), in aneuploid cells and other G1 arrests. This analysis revealed that the composition of SASP varied between different G1 arrests (Figure 3E, S2C). However, ArCK and G1 arrested cells secreted a plethora of chemokines and cytokines including factors contributing to NK cell recognition (*e.g*., CCL2 and MIF), whereas aneuploid cycling cells did not. We conclude that ArCK cells attract NK cells at least in part by expressing a classic senescence immune recognition program. However, aneuploid cycling cells, which are recognized almost as well as ArCK cells do not. What causes their recognition is not known. If it involves secreted factors, they are not included in the chemokine and cytokine panel that we tested.

### Cell autonomous and non-cell autonomous contribute to NK cell-mediated cytotoxicity towards aneuploid cells

Aneuploid cycling cells attracted NK cells by mechanisms other than inducing a senescence program. To determine how this occurred, we first asked whether this recognition required secreted factors. We collected the culture medium from aneuploid cells and applied it to euploid cells (Figure 4A and B). Medium previously used to culture ArCK or aneuploid cycling cells for 12 hours increased NK-mediated cytotoxicity by ~1.5 fold towards euploid cells.

**Figure 4.**
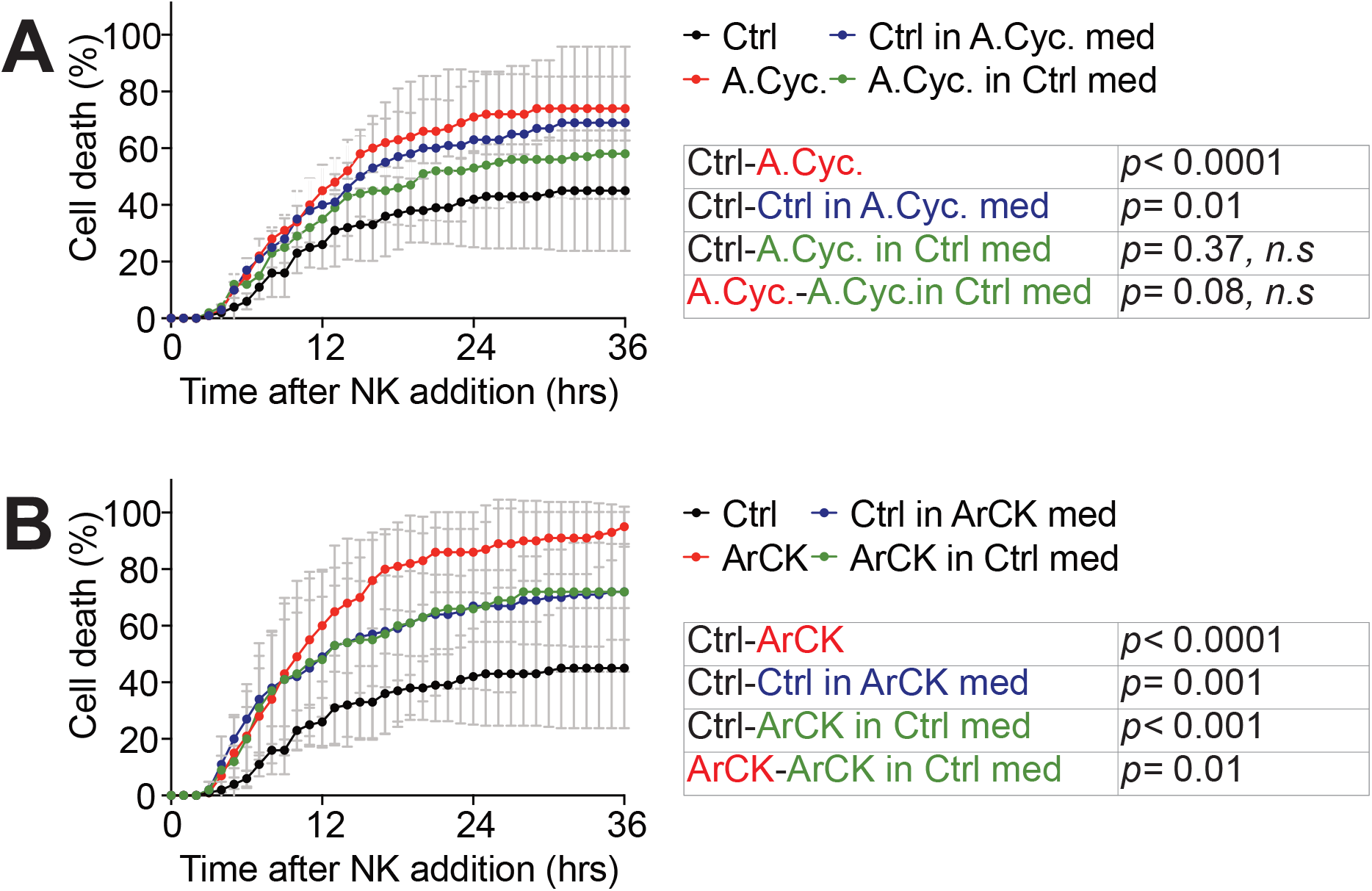
Secreted factors are primarily responsible for NK cell mediated-elimination of aneuploid cells. (A) NK cell medium was incubated with either euploid or aneuploid cycling cells for 12 hours. At the time of NK cell addition, media were switched between aneuploid cycling cells (A. Cyc.) and euploid control cells (Ctrl). NK cell killing was measured as described in Figure 1B. For reference, NK cell killing experiments of aneuploid cycling (A.Cyc) and euploid control cells (Ctrl) without medium switch were performed side by side and plotted on the graph. Black, euploid control cells without medium switching; red, aneuploid cycling cells without medium switching; blue, euploid control cells in aneuploid cycling cell condition medium; green, aneuploid cycling cells in euploid control cell condition medium (n=2). Statistical analyses were performed as in Figure 1B (*n.s* – not significant). (B) NK cell medium was incubated with either euploid or ArCK cells for 12 hours and the experiment was performed as described in (A). Black, euploid control cells without medium switching; red, ArCK cells without medium switching; blue, euploid control cells in ArCK cell condition medium; green, ArCK cells in euploid control cell condition medium (n=2). Statistical analyses were performed as in Figure 1B.

When we used pre-conditioned medium from euploid cells to co-culture NK cells with ArCK cells, we observed a decrease in NK-mediated cytotoxicity towards these cells (Figure 4B). However, conditioned medium switch did not completely abolish the differences in NK cell-mediated killing between aneuploid cells and their euploid counter parts. ArCK cells still were killed more by NK cells than euploid cells when co-cultured in preconditioned medium from euploid cells (Figure 4B). This could be due to the accumulation of secreted factors during the 36-hour live cell imaging as well as ArCK cell surface features that mediate their elimination by NK cells.

When we used pre-conditioned medium from euploid cells to co-culture NK cells with aneuploid cycling cells we found that NK-mediated cytotoxicity towards these cells was largely mediated by secreted factors (Figure 4A). There continued to be a subtle difference in NK cell-mediated killing between aneuploid cycling cells and their euploid counter parts but this difference was merely a trend and not statistically significant (Figure 4A). We conclude that NK-cell mediated killing of aneuploid cells is largely mediated by secreted factors, but cell surface signals also contribute, especially in the case of ArCK cells.

### NF-κB and interferon-mediated pathways are upregulated in aneuploid cells

Having established that both secreted factors and cell surface features of aneuploid cells contribute to NK cell-mediated killing, we next wished to determine how NK cell recognition is induced. For this, we profiled the gene expression of ArCK cells and aneuploid cycling cells and compared it to doxorubicin, palbociclib or nutlin3-treated cells. Hierarchical clustering of the differential expression data indicated that all cell-cycle arrested samples (ArCK and doxorubicin, palbociclib or nutlin3-treated cells) clustered together whereas the aneuploid cycling cells were closer to the euploid untreated control cells (Figure S3A). Gene set enrichment analysis (GSEA) revealed common hallmarks that were significantly upregulated across all conditions, such as protein secretion and the p53 pathway. The hallmark gene set “TNFalpha–via NF-κB signaling” was among the most upregulated pathways in aneuploid and doxorubicin-treated samples (Figure S3B, 5A and B). ArCK cells also exhibited increased expression of the interferon alpha and gamma response, immune complement, JAK-STAT, and interleukin (IL) related pathways (Figure 5A). In palbociclib or nutlin3-treated samples, we observed only a mild upregulation of immune related signatures, but none of them reached significance (*p* < 0.05).

**Figure 5.**
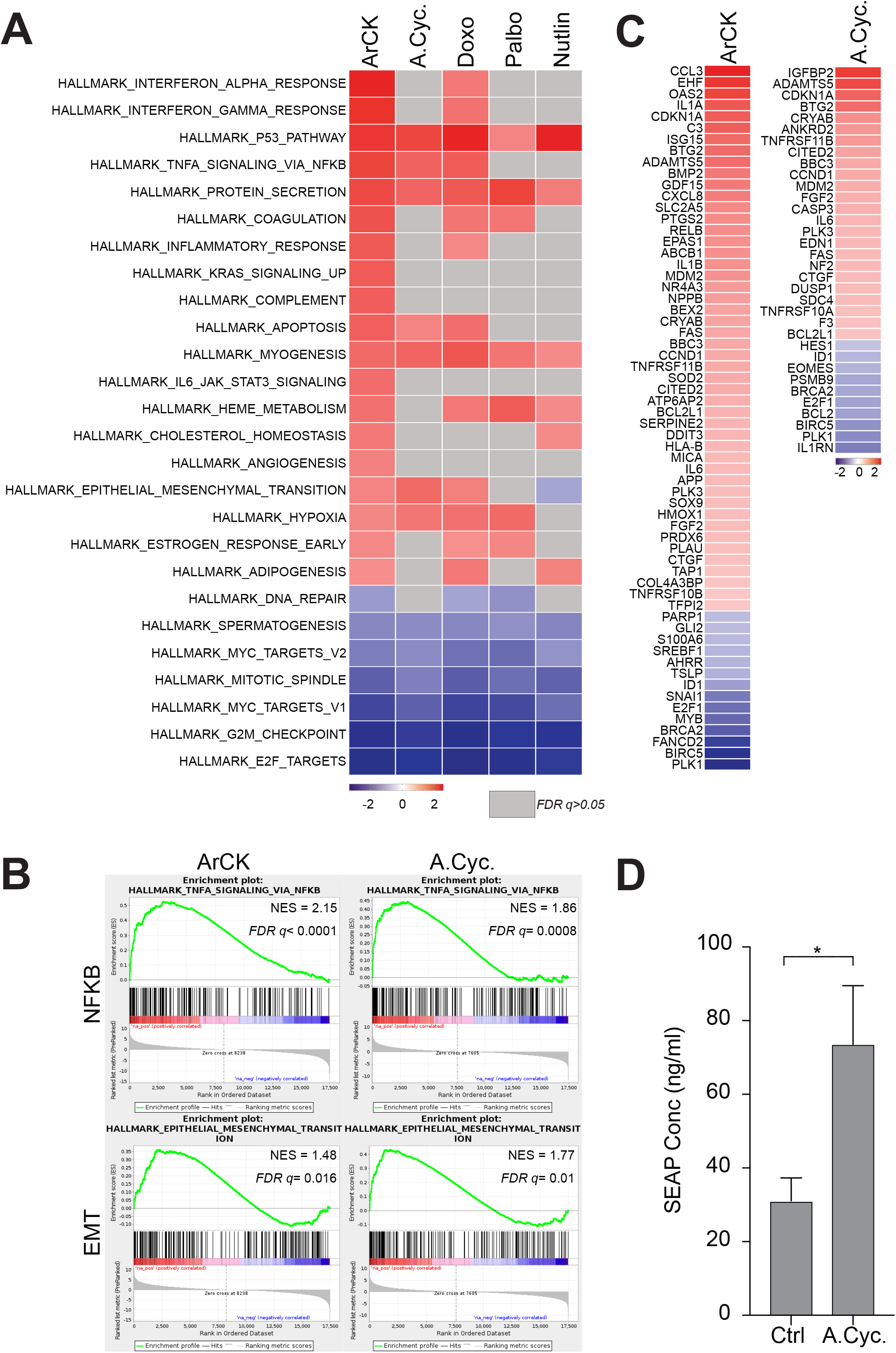
NF-κB pathway activation in aneuploid cells. (A) Significantly differentially expressed hallmarks in ArCK cells compared to euploid control cells are shown in the first column in the heatmap. Normalized enrichment scores were plotted (*p* value < 0.05). The corresponding NES for these hallmarks in aneuploid cycling (A.Cyc.), doxorubicin (Doxo), palbociclib (Palbo), and nutlin 3 (Nutlin) treated cells were also plotted. Hallmarks did not reach statistical significance (*p* > 0.05) were shown in grey. (B) Enrichment plots for “TNFalpha signaling via NFKB” and “Epithelial_mesenchymal_transition” hallmark signatures in ArCK and aneuploid cycling (A.Cyc.) cells compared to the euploid proliferating counterparts. (C) Significantly differentially expressed RelA and RelB target genes in ArCK cells and aneuploid cycling cells (A.Cyc.) compared to euploid control cells (Log2 fold change, *p* value < 0.05). (D) Measurement of NF-κB activity using a NF-κB alkaline phosphatase (SEAP) reporter assay. The reporter was expressed in RPE1-hTERT cells by transient transfection. 10-12 hours after transfection, cells were treated with either DMSO (Ctrl) or reversine (500 nM) for 60 hours (aneuploid cycling, A.Cyc.) and the secretion of alkaline phosphatase in the culture supernatant was measured (two-tailed t-test, *p*= 0.049).

Given the importance of NF-κB in mediating immune recognition, we characterized this pathway further in aneuploid cycling and ArCK cells. RNA-seq analysis revealed significant transcriptional upregulation of both RelA and RelB target genes in ArCK cells and in aneuploid cycling cells, although up-regulation was less prominent in the latter (Figure 5C). To substantiate the finding that NF-κB was activated in aneuploid cycling populations, we employed an assay in which the expression and secretion of alkaline phosphatase (AP) is controlled by an NF-κB regulatory element (Signorino et al. 2014). Using this assay, we observed a ~2 fold increase in AP secretion in aneuploid cycling cells (Figure 5D and S4A). We however observed neither accumulation of NF-κB in the nucleus nor IκB phosphorylation (Figure S4B and C) suggesting that the pathway may only be transiently active or at a low level. We conclude that the NF-κB pathway is activated in both ArCK and aneuploid cycling cells.

### The canonical and non-canonical NF-κB pathway is required for NK cell-mediated killing of aneuploid cells

What is the relevance of the NF-κB pathway in NK cell-mediated elimination of aneuploid cells? To address this question, we generated *RELA* and *RELB* single KO cell lines using CRISPR-Cas9 (Figure S5A and B). Inactivation of either transcription factor did not significantly affect NK-mediated cytotoxicity towards ArCK and aneuploid cycling cells in most of the KO clones (Figure 6A and B, Figure S5C and D). However, when we inactivated both *RELA* and *RELB*, NK cell-mediated killing in ArCK was significantly reduced to a level comparable to the killing of proliferating euploid controls (Figure 6C and S5E). Similar results were observed when aneuploid cycling and ArCK cells were treated with a pan-NF-κB inhibitor that interferes with both catalytic subunits of IKK (with different binding affinities; (Yang et al. 2006)) to block canonical and alternative NF-κB activation (Figure 6D). These data indicate that aneuploidy triggers NF-κB activation, which contributes to their recognition and elimination by NK cells.

**Figure 6.**
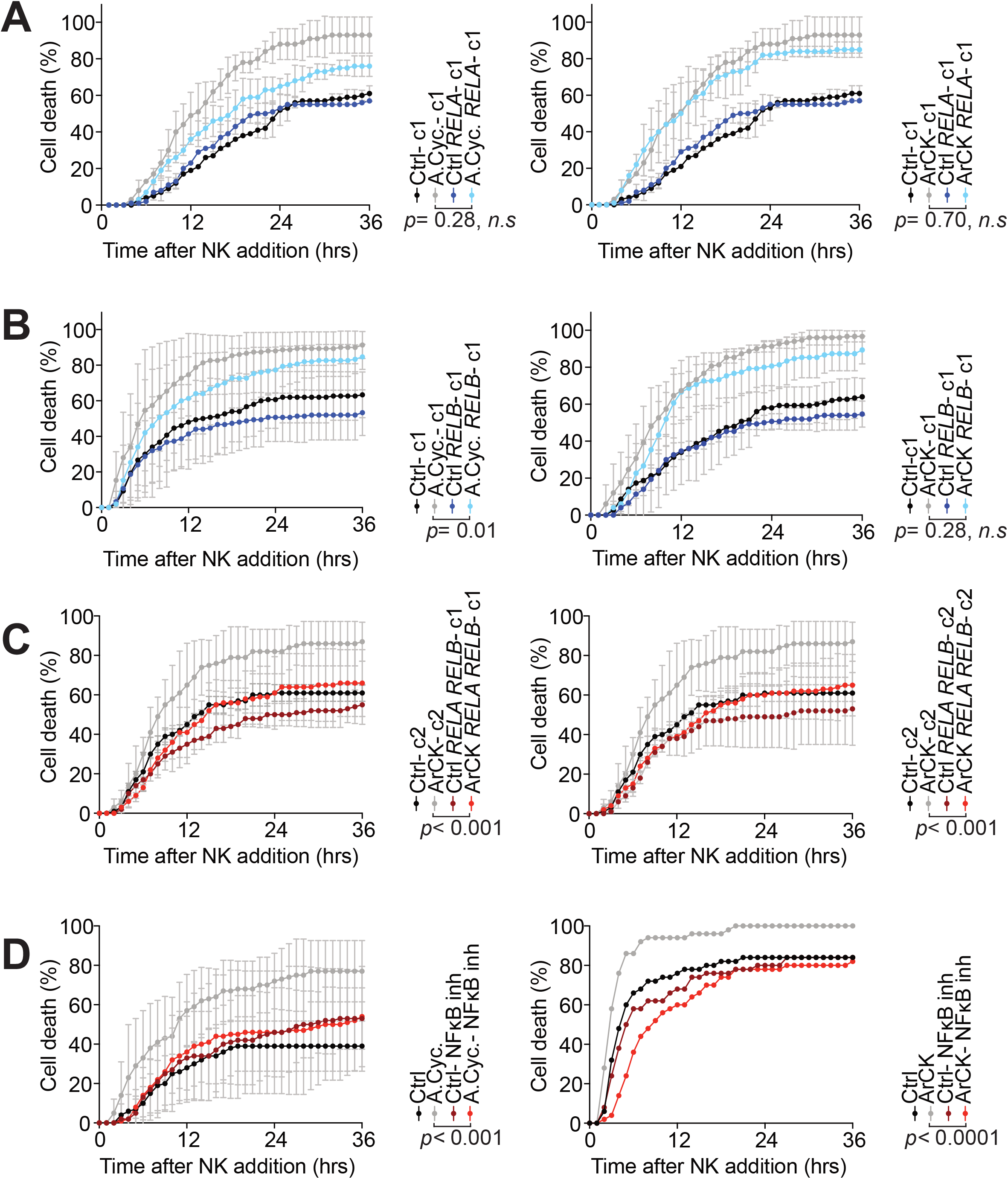
The canonical and non-canonical NF-κB pathway are required for NK cell-mediated killing of aneuploid cells. (A) *RELA* KO RPE1-hTERT cell lines were generated using the CRISPR-Cas9 method and single cell clones were isolated. Aneuploid cycling (A.Cyc.) and ArCK *RELA* knock out cells were generated as described in Figure 1A and 1E. NK cell-mediated cell death in each condition was measured as described in Figure 1B. The NK cell killing of *RELA* aneuploid cycling (A.Cyc.) and ArCK cells was compared to single cell clones that harbor an empty vector (n=2; mean ± SD). Statistical analyses were performed as in Figure 1B (*n.s* – not significant). (B) The effect of inactivating RelB on NK cell-mediated cytotoxicity in aneuploid cells. The same experimental methods were used as described in (A). (C) *RELA, RELB* double KO RPE1-hTERT cells were generated using the CRISPR-Cas9 method and single cell clones were obtained. ArCK *RELA, RELB* KO cells were generated as described in Figure 1A. The NK cell killing of ArCK *RELA, RELB* KO cells was compared to single cell clones that harbor empty vectors (n=2; mean ± SD). Statistical analyses were performed as in Figure 1B. (D) Aneuploid cycling (A.Cyc.) and ArCK cells were generated as described in Figure 1. Aneuploid cycling, ArCK or euploid proliferating cells were treated with either DMSO or a pan-NF-κB inhibitor BMS-345541 (5 μM) for 48 hours before assessing NK cell-mediated cytotoxicity. The drug was washed out during the NK cell co-culture assay. Statistical analyses were performed as in Figure 1B.

ArCK cells also induce interferon alpha and gamma responses as judged by RNA-seq and RT-qPCR (Figure S6A and B). The alpha and gamma interferon responses are primarily mediated by the JAK-STAT pathway. We confirmed that activation of these two gene expression signatures was indeed mediated by the JAK-STAT pathway as inactivation of *STAT1* by CRISPR-Cas9 reduced both the interferon alpha and interferon gamma responses in ArCK cells (Figure S6C and D). To determine the biological relevance of this JAK-STAT response we depleted *STAT1* in both aneuploid cycling and ArCK cells. Knock-down of *STAT1* did not significantly affect the ability of NK cells to kill ArCK and aneuploid cycling cells in most of the clones (Figure S7). Our data indicate that *STAT1* is involved in potentiating interferon secretion which may facilitate immune recognition of aneuploid cells. Deletion of *STAT1* alone is not sufficient, however, to significantly affect NK cell-mediated cell death of aneuploid cells. The NF-κB pathway appears to be the major mediator of this immunogenicity.

### NF-κB is active in highly aneuploid cancer cell lines

High levels of aneuploidy correlates with immune evasion in cancer (Davoli et al. 2017; Taylor et al. 2018). Our data indicate that in untransformed cells, aneuploidy is associated with an NF-κB response that is relevant for the recognition of aneuploid cells by NK cells *in vitro*. It is thus possible that the transformed state of cancer cells suppressed aneuploidy-induced NF-κB signaling. To test this hypothesis, we interrogated the association between NF-κB activation and the degree of aneuploidy across the cancer cell line encyclopedia (CCLE; (Barretina et al. 2012; Ghandi et al. 2019)). Degree of aneuploidy was scored in the almost 1000 cell lines in the CCLE as described in Cohen-Sharir et al. (in revision). We then created two groups of cell lines, an aneuploid and a near-euploid group, defined as the top and bottom quartiles of the number of arm-level chromosome gains and losses, respectively. To assess NF-κB activity in these two cell line groups, we created a ssGSEA signature (Subramanian et al. 2005) score for the Hallmark_TNFA_signaling_via_NFKB gene set and computed the association between this signature and degree of aneuploidy by linear regression analysis (see Methods). In agreement with our hypothesis, we found that highly aneuploid cancer cell lines show a higher gene expression levels of NF-κB activation compared to lowly aneuploid lines (Figure 7). These results indicate that an NF-κB response is evident even in highly aneuploid cancer cell lines and that loss of NF-κB signaling is not the cause of immune evasion in highly aneuploid cancer cell lines.

**Figure 7.**
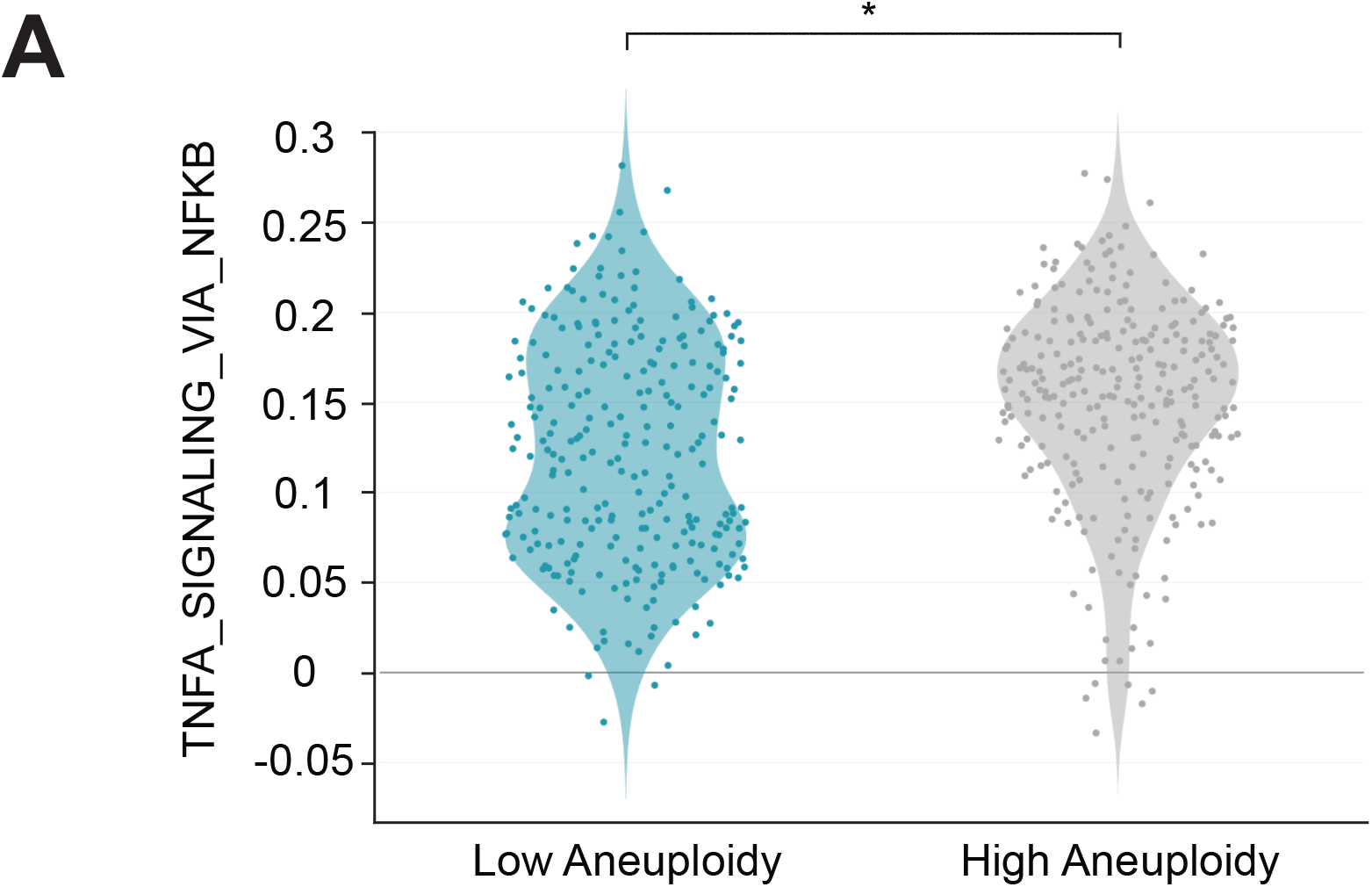
NF-κB is active in highly aneuploid cancer cell lines. (A) Comparison of the Hallmark gene expression signature “TNFA_signaling via NFKB” in neardiploid (low aneuploidy) and highly-aneuploid (high aneuploidy) human cancer cell lines from the CCLE. *, *p*= 5e-08, empirical Bayes-moderated t-statistics. The y-axis represents the ssGSEA expression score.

## Discussion

We previously found that untransformed cells that underwent senescence due to highly abnormal karyotypes are recognized by NK cells *in vitro*. Here we show that aneuploid cells that are still dividing can also be recognized and eliminated by NK cells *in vitro*. Our data further indicate that NF-κB signaling is central to this interaction between aneuploid cells and NK cells.

### The NF-κB pathway induces immunogenicity of aneuploid cells

Multiple studies have shown that senescent cells are recognized and eliminated by NK cells *in vitro*. In this study, we investigated how aneuploidy-induced senescence causes NK cell recognition. We first tested the hypothesis that G1 arrest elicits an NK cell response by comparing NK cell killing kinetics between various G1 arrests. This analysis revealed that G1 arrest *per se* is not sufficient to cause recognition by NK cells. mTOR inhibition, which causes a quiescence-like G1 arrest, did not elicit an NK cell response. Instead, our studies indicate that as previously described for other NK cell – senescent cell interactions, the senescence-associated secretory program is primarily responsible for NK cell recognition of aneuploid cells (Iannello et al. 2013). Medium harvested from aneuploid cell cultures increases the ability of NK cells to kill euploid cells. Even a small number of aneuploid cells in a culture can induce wide-spread NK cell killing of euploid cells (unpublished observations), suggesting that aneuploid cells can establish a pro-inflammatory environment where immune clearance of not just aneuploid cells is taking place. The senescence-associated secretary program is, however, not the only route whereby aneuploid cells elicit NK cell recognition. We find that aneuploid cycling cells are more effectively recognized and killed by NK cells than euploid cells even though they do not exhibit a senescence-associated secretory program.

Our data indicate that the major inducer of immunogenicity of aneuploid cells is the NF-κB pathway. Inactivation of the canonical and non-canonical NF-κB pathways is sufficient to protect aneuploid cells from NK cell-mediated killing *in vitro*. However, we note that the degree of NF-κB activation observed in aneuploid cells is low compared to what is observed in response to acute stimuli such as TNF-α. Furthermore, although NF-κB target genes are significantly upregulated in aneuploid cells, IκB phosphorylation, a measure for NF-κB pathway activity, was not evident in aneuploid cells. This observation suggests that NF-κB activation in aneuploid cells occurs at a low level but does so continuously creating a chronic inflammatory environment.

Could other immune-response inducing pathways contribute to aneuploid cell recognition by NK cells? Chromosome mis-segregation has been linked to induction of an interferon response (Bakhoum et al. 2018; Vasudevan et al. 2020). We do observe up-regulation of the alpha and gamma interferon response in ArCK cells but not in aneuploid cycling cells. Our data further indicate that inactivation of *STAT1*, the major transcription factor mediating the interferon response, does not significantly affect elimination of aneuploid cells by NK cells. *STAT1* activation has been observed in untransformed and cancer cell lines where aneuploidy was induced by continuous exposure to reversine (F. Foijer, personal communication). In this experimental set up, persistent DNA damage, which accompanies CIN rather than aneuploidy *perse* is likely to be the primary cause for JAK/STAT1 pathway activation. In aneuploid cells that do not experience CIN the pathway does not significantly contribute to immunogenicity.

Chromosome mis-segregation also generates micronuclei (S. Liu et al. 2018). The nuclear envelope of these micronuclei is unstable leading to their frequent rupture. This in turn causes DNA to spill into the cytoplasm, which activates the cGAS-STING pathway (Dou et al. 2017). cGAS-STING then causes induction of the NF-κB and interferon response (Dunphy et al. 2018). In our experimental set up, over 50% of aneuploid cycling cells harbor micronuclei, yet cGAS-STING pathway activation was, at best, weak as judged by the low degree of IRF3 phosphorylation (Figure S8A). Furthermore, STING inactivation did not affect NK cell-mediated killing of aneuploid cells (Figure S8B). Together, these data indicate that at least in our *in vitro* assay, NF-κB is the predominant pathway by which aneuploidy induces NK cell-mediated cytotoxicity.

### Multiple signals activate NF-κB in aneuploid cells

A key question arising from our findings is what causes NF-κB activation in aneuploid cells. Micronuclei do not appear to be a major source of immune pathway activation. Retrotransposon activation, which occurs in response to several cellular stresses including senescence, may however be a cause. We found that in aneuploid cells, two dsRNA sensors, Rig-I (*DDX58*) and Mda5 (*IFIH1*) are induced (Figure S8C). Furthermore, ORF1p, one of the proteins encoded by the LINE-1 retrotransposon, was mildly induced in ArCK cells (Fig S8D). This upregulation of retrotransposons could be relevant to NK cell recognition of aneuploid cells. Inhibition of reverse transcription, by treating cells with the cytosine analog 2’,3’-dideoxy-3’-thiacytidine (3TC) partially suppressed NK cell-mediated cytotoxicity towards aneuploid cells (Figure S8E).

While increased retrotransposition could contribute to NF-κB activation in aneuploid cells, the aneuploidy-associated stresses are likely to be the major activators of this immune-response inducing pathway. We previously found that the surface molecules MICA, MICB, CD155, CD112, ULBP1 and ULBP2-that mediate NK cell recognition-are subtly (about 2-fold) upregulated in ArCK cells (Santaguida et al. 2017). The increased surface presentation of these molecules was the result of specific cellular stresses. In particular, MICA and MICB are activated in response to proteotoxic stress. CD112 (also known as Nectin-2) and CD155 (also known as PVR), are expressed in response to DNA damage. ULBP1 and ULBP2 are the product of cellular stresses and DNA damage (Raulet & Guerra 2009). Proteotoxic, oxidative and genotoxic stress are defining features of the aneuploid state (reviewed in (Santaguida & Amon 2015)). The observation that immunogenic cell surface molecules that are induced in response to these stresses are presented at increased levels at the surface of aneuploid cells indicates that multiple features of the aneuploid state contribute to immunogenicity. We propose that aneuploid cells do not express a unique NK cell attracting feature but rather a combination of stresses elicited by the aneuploid state mediate the interaction between aneuploid cells and NK cells.

### Immune recognition of aneuploidy in cancer

Aneuploidy is a hallmark of cancer that correlates with aggressive disease and immune evasion. Yet in primary cells there is ample evidence that chromosome instability as well as aneuploidy elicit a variety of immune responses. Micronuclei activate the interferon response via the cGAS/STING pathway (Mackenzie et al. 2017; Dou et al. 2017) and aneuploid cells become immunogenic through activation of the NF-κB pathway. Chromosome instability upregulates stress-activated protein kinase (SAPK) and c-Jun N-terminal kinase (JNK) pathways which contribute to inflammatory response (Clemente-Ruiz et al. 2016; Benhra et al. 2018). MHC complex and antigen processing gene signatures have also been shown to be associated with aneuploidy (Dürrbaum et al. 2014). Hence, a crucial question in the field, is how malignant transformation dampens aneuploidy and CIN-induced immunogenicity. One possibility we investigated was that NF-κB signaling was down-regulated in highly aneuploid cancer cell lines. This was not the case-in fact the opposite appeared to be true. The more aneuploid the cell line, the higher NF-κB signaling levels were. Aneuploidy associated stresses could still exist in highly aneuploid cancer cell lines. It is also possible that high level of NF-κB signaling is preferentially selected in highly aneuploid cancer cell lines because NF-κB antagonizes p53, which is known to inhibit the growth of highly aneuploid cells (M. Li et al. 2010; Rahnamoun et al. 2017). Irrespective of why NF-κB signaling is upregulated in highly aneuploid cancer cell lines, immune signaling has not been silenced. However, in tumor samples, the degree of NF-κB activation inversely correlates with degree of aneuploidy (Taylor et al. 2018). These results raise the interesting possibility that silencing of aneuploidy-induced immunogenicity is a non-cell autonomous event in cancer, perhaps induced by cells in the tumor’s microenvironment. Understanding which aspects of aneuploidy activate NF-κB signaling and how signaling activity of the pathway is modulated during tumor evolution will be the critical next steps in understanding the role of aneuploidy during tumorigenesis.

## Author contributions

Conceptualization, R.W.W., A.A. and S.S.; Investigation, R.W.W., S.V., U.B.-D. and S.S.; Writing, R.W.W., U.B.-D., A.A. and S.S.; Funding Acquisition and Supervision, U.B.-D., A.A. and S.S. All authors discussed the results and commented on the manuscript.

## Acknowledgements

We thank Florijs Foijer and members of the Amon and Santaguida labs for their helpful comments and discussions regarding this project and manuscript. We thank Charlie Whittaker, Dikshant Pradhan, and the Swanson Biotechnology Center for help with the gene expression analysis. This work was supported by NIH grant CA206157 to A.A., who is an investigator of the Howard Hughes Medical Institute, the Paul F. Glenn Center for Biology of Aging Research at MIT and the Ludwig Center at MIT and by grants to S. S. from the Italian Association for Cancer Research (MFAG 2018 - ID. 21665 project), Fondazione Cariplo, Ricerca Finalizzata (GR-2018-12367077), the Rita-Levi Montalcini program from MIUR and by the Italian Ministry of Health with Ricerca Corrente and 5×1000 funds. Work in the Ben-David lab is supported by the Azrieli Foundation, the Richard Eimert Research Fund on Solid Tumors, the Tel-Aviv University Cancer Biology Research Center, and the Israel Cancer Association.

---

**Figure S1.**
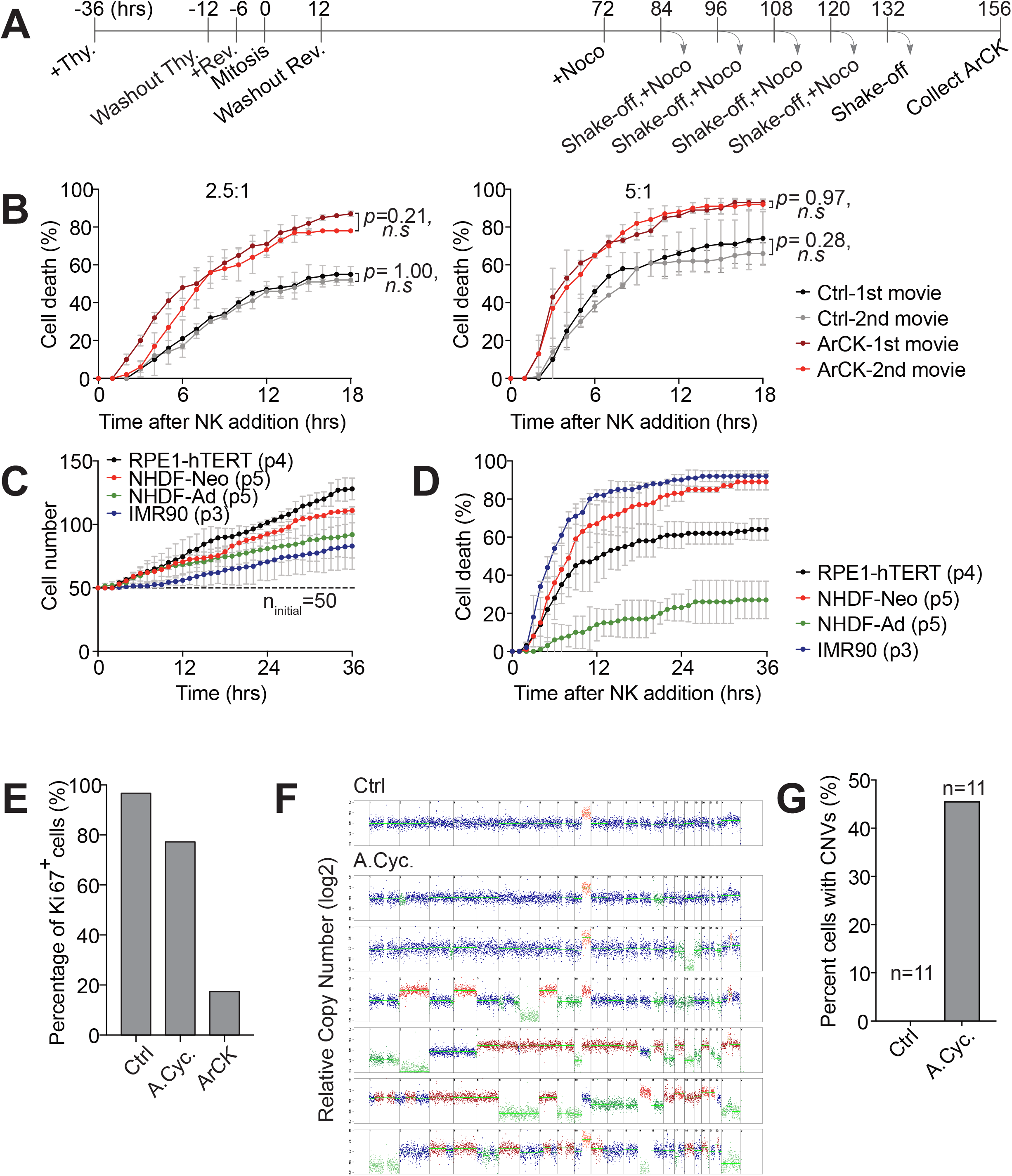
Characterization of the NK cell killing assay. (A) Experimental scheme for the generation of ArCK populations. The timeline is relative to the estimated reversine-induced chromosome mis-segregation event (0h). (B) Measurement of NK cell-mediated killing of ArCK cells in two consecutive 18h time lapse experiments. Cells were cultured as described in Figure 1A and were analyzed by time lapse microscopy. After the first 18 hours of the analysis, the cell suspension was collected and co-cultured with a second set of target cells. NK cell-mediated killing was measured in the first (black and dark red curves) and the second (grey and light red curves) 18h time lapse and plotted on the same graph. The killing assay was performed at a NK cell to target cell ratio of 2.5:1 (left panel) and 5:1 (right panel; n=2; mean ± SD). Statistical analyses were performed as in Figure 1B (*n.s*-not significant). (C) Cell proliferation measurements in the absence of NK cells. RPE1-hTERT (passage 4), human normal neonatal or adult human dermal fibroblasts (NHDF-Neo, passage 5 or NHDF-Ad, passage 5), and human embryonic lung fibroblast (IMR90, passage 3) were plated side by side in NK cell medium and cell proliferation rate was recorded using live cell imaging as described in Figure 1C. The dashed line indicates the starting cell number (ninitial=50; n=2; mean ± SD). (D) NK cell-mediated cytotoxicity across different cell types. The killing of RPE1-hTERT, human normal neonatal or adult human dermal fibroblasts (NHDF-Neo or NHDF-Ad), and human embryonic lung fibroblast (IMR90) were measured as described in Figure 1B using a NK cell to target cell ratio of 2.5 to 1 (n=2; mean ± SD). (E) The percentage of Ki67 positive cells in aneuploid cell population. ArCK, aneuploid cycling cells (A.Cyc.), and euploid cells (Ctrl) were cultured as described in Figure 1A and 1E. Ki67 staining was evaluated at the time corresponding to the end of NK cell killing assay (n ≥ 100 cells per condition). (F) Segmentation plots of control (Ctrl) and aneuploid cycling (A. Cyc.) cells based on single cell DNA sequencing. Copy number of single cells relative to euploid reference is shown (n=11). Representative plots from the two populations are shown. A clonal chromosome 10q gain is observed across the population. (G) Quantification of percentage of cells with copy number variations (CNVs). The pre-existing gain of chromosome 10q was not counted in the analysis.

**Figure S2.**
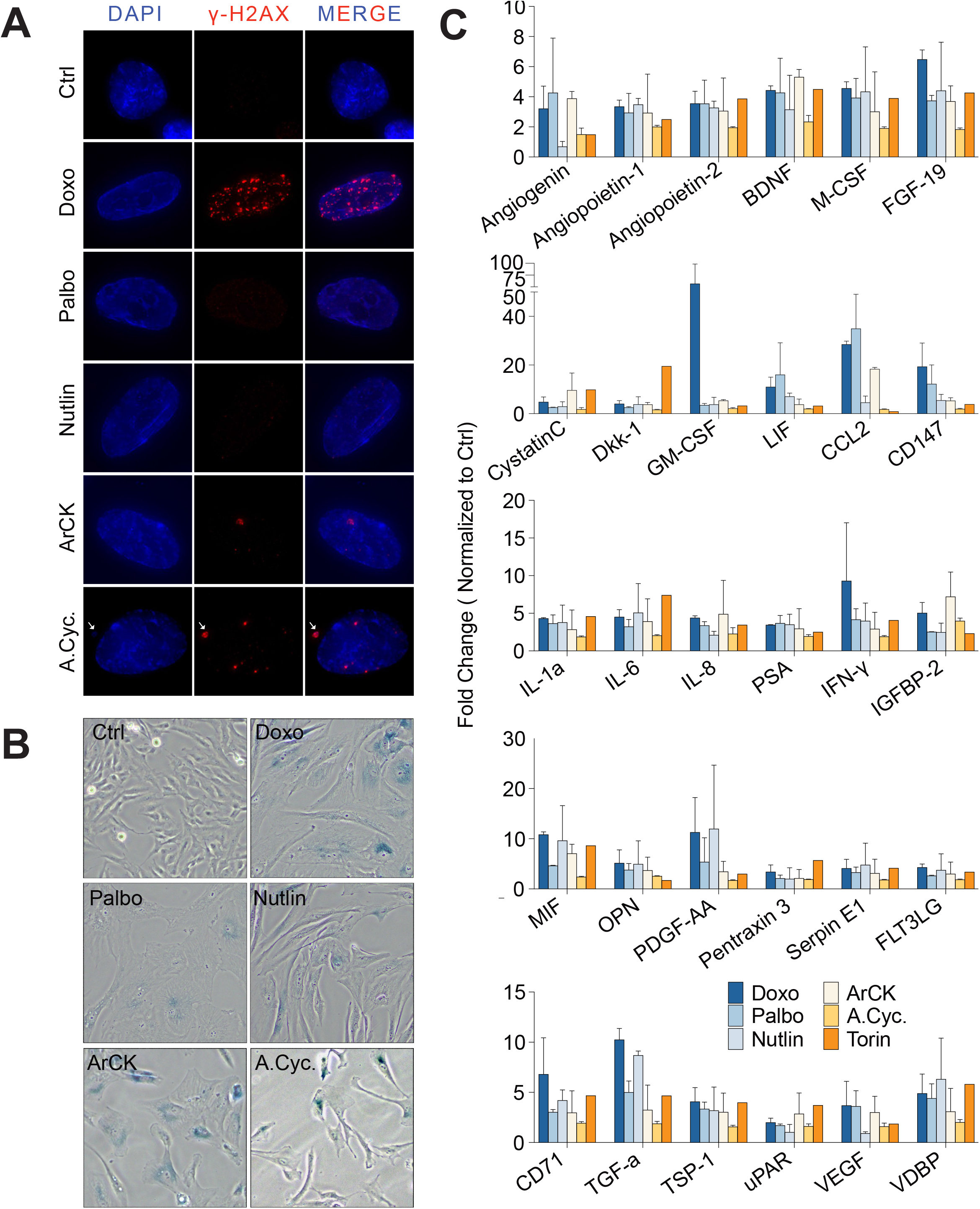
Senescence phenotypes in aneuploid and G1-arrested cells. (A) Representative images of g-H2AX staining in the indicated samples. g-H2AX is in red and DNA in blue. A micronucleus in aneuploid cycling cells (A.Cyc.) is highlighted by a white arrow. (B) Representative image of senescence-associated β-galactosidase staining in the indicated samples. (C) Analysis of all cytokines secreted by the indicated cells. Cytokine levels are shown as fold change of euploid control cells (mean ± SD).

**Figure S3.**
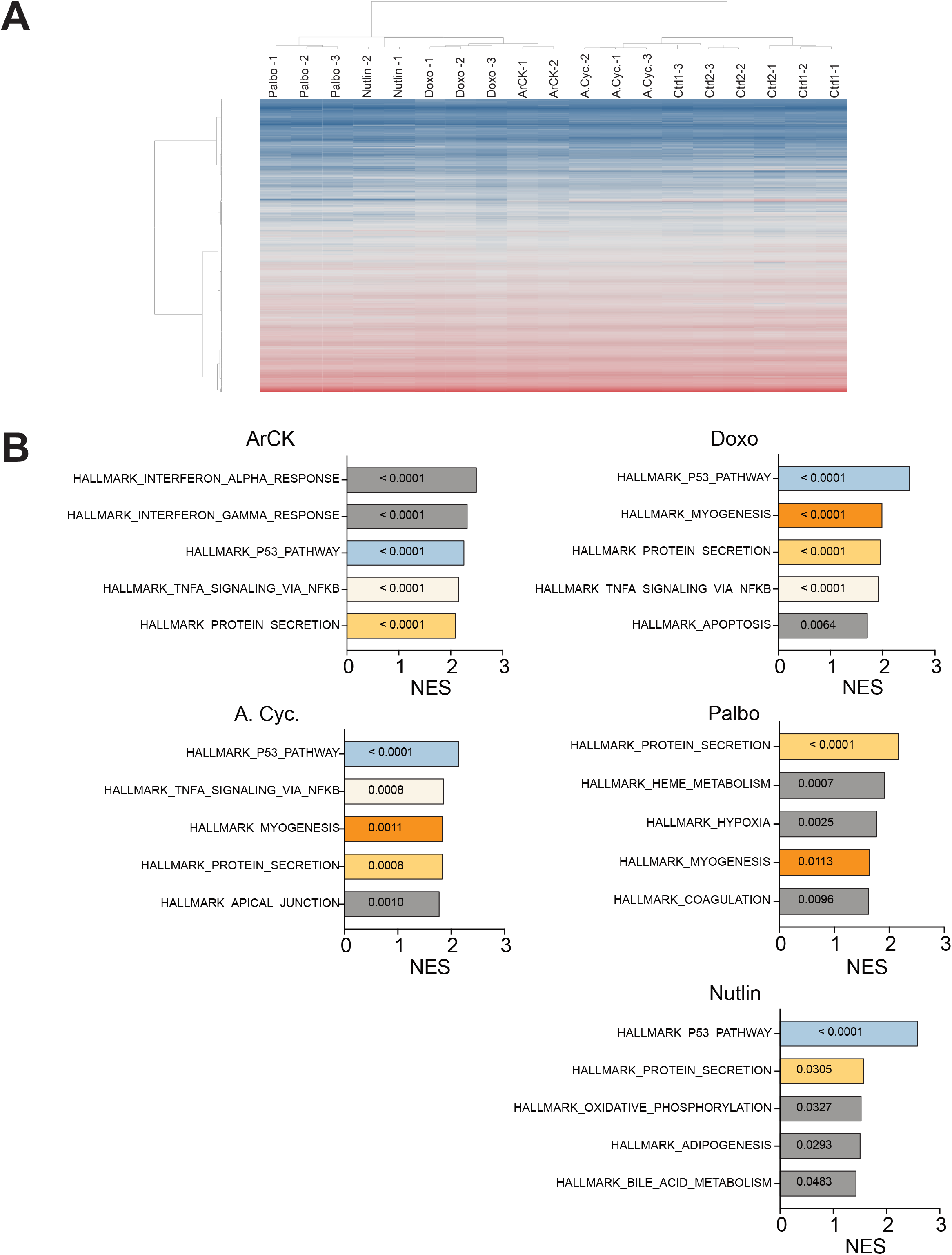
Gene expression analysis in aneuploid cells and G1-arrested cells. (A) Hierarchical clustering of differential RNA expression analysis of euploid proliferating control (Ctrl), doxorubicin (Doxo), palbociclib (Palbo), nutlin3 (Nutlin)-treated, ArCK, and aneuploid cycling (A.Cyc.) samples. The samples were generated as described in Figure 1 and 2. (B) Gene set enrichment analysis (GSEA) for doxorubicin (Doxo), palbociclib (Palbo), nutlin 3 (Nutlin)-treated, ArCK, and aneuploid cycling (A.Cyc.) cells relative to euploid proliferating control cells. The normalized enrichment score (NES) for the top five upregulated hallmarks in each condition were plotted. The numbers on the NES score bar indicated the corresponding *p-*values for each hallmark.

**Figure S4.**
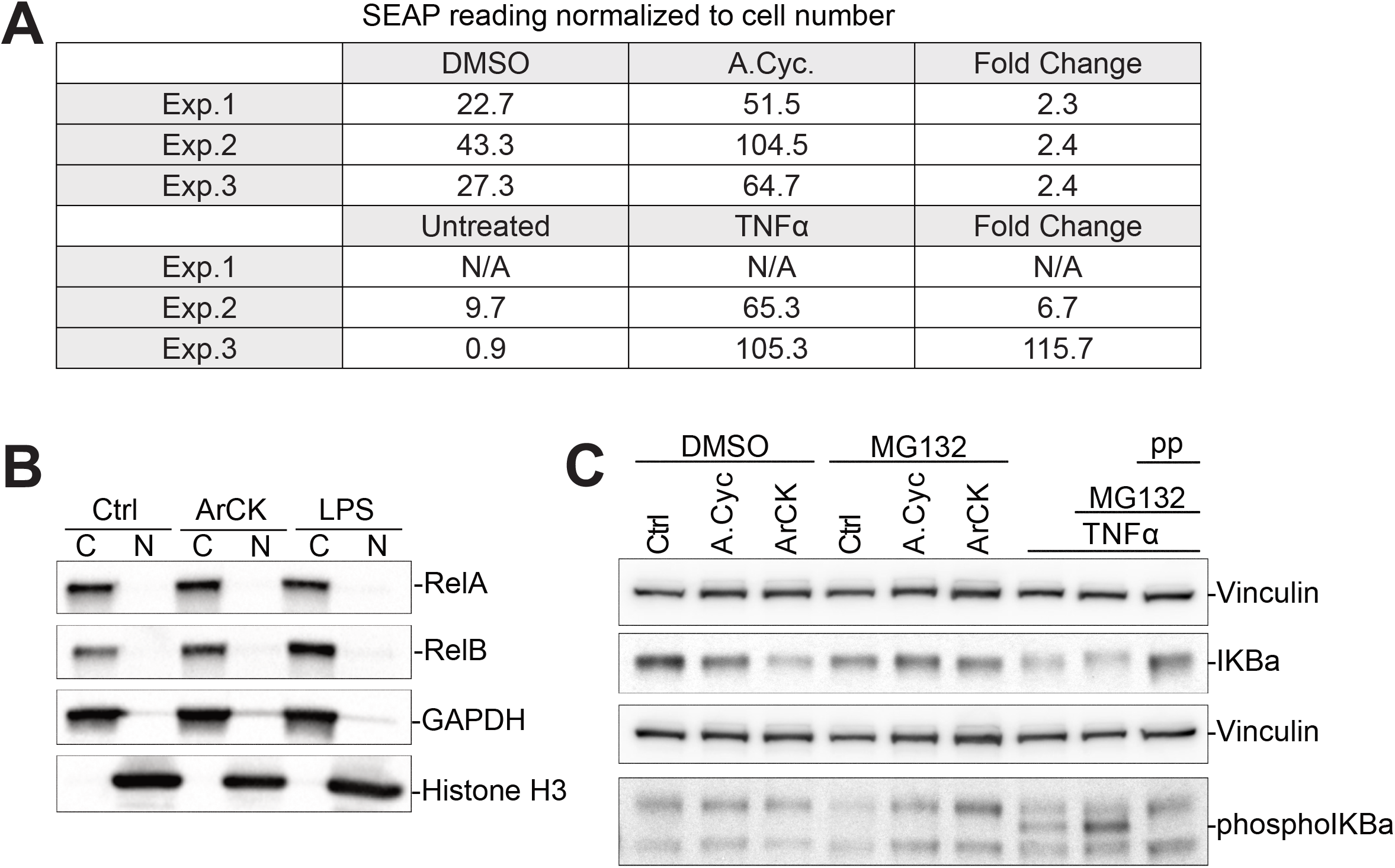
NF-κB pathway activity in aneuploid cells. (A) NF-κB alkaline phosphatase (SEAP) reporter absorbance readings normalized to cell number for individual experiment. As a positive control, RPE1-hTERT cells were treated with TNFα (30 ng/ml) for 24 hours. The fold change for aneuploid condition was calculated as aneuploid cycling (A.Cyc.)/DMSO. For the positive control, the fold change was calculated as TNF-α/untreated. (B) Measurement of RelA or RelB nuclear translocation. ArCK cells were generated as described in Figure 1A. LPS (1 ng/mL) treatment was used as a positive control. GAPDH is a cytoplasmic protein, histone H3 is a nuclear protein. They served as controls for effective fractionation (n=2; results were comparable). (C) Aneuploid cycling (A.Cyc.) cells and ArCK cells were generated as described in Figure 1. Proteasome inhibitor MG132 (1 μm) was applied 1.5 hours prior to protein extraction as indicated (lanes 4-6 and lanes 8-9). We treated cells with MG132 to detect low levels of phosphorylated IκB, which gets degraded upon phosphorylation (Perkins 2007). As a positive control, RPE1-hTERT cells were stimulated with TNF-α (10 ng/mL) for one hour (lanes 7-9). To confirm the specificity of the phospho epitope, protein lysates from TNF-α stimulated cells were treated with lambda phosphatase (lane 9).

**Figure S5.**
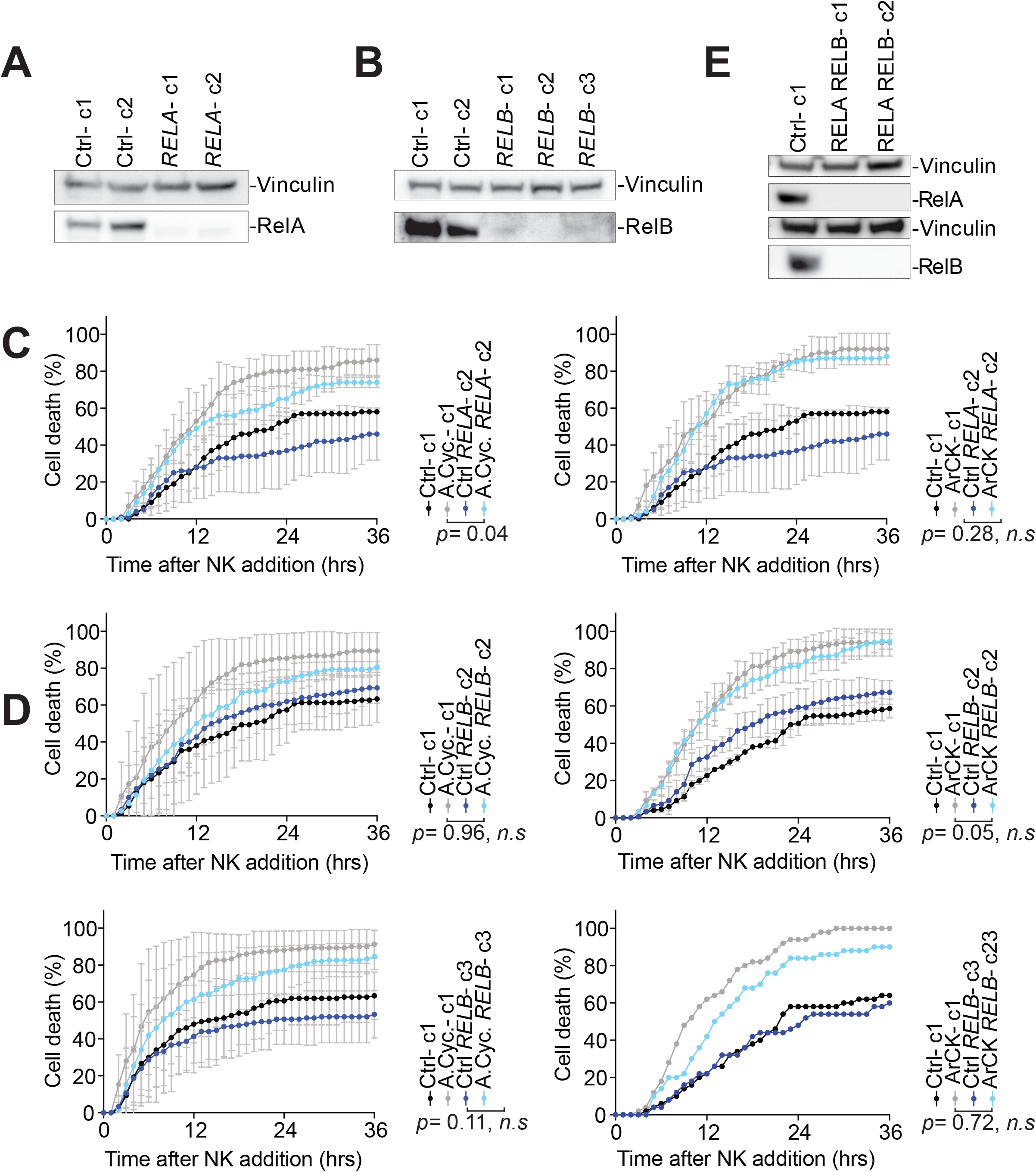
Effects of RelA and RelB inactivation on the immunogenicity of aneuploid cells. (A) Measurement of RelA protein levels in *RELA* KO single cell clones generated in RPE1-hTERT cells. Vinculin was used as a loading control. (B) Measurement of RelB protein levels in *RELB* KO single cell clones generated in RPE1-hTERT cells. (C) The effect of inactivating RelA on NK cell-mediated cytotoxicity in aneuploid cells. The experiment was performed as described in Figure 6A, except a different *RELA* KO single cell clone was used. (D) The effect of inactivating RelB on NK cell-mediated cytotoxicity in aneuploid cells. The experiment was performed as described in Figure 6B, except a different *RELB* KO single cell clones was used. (E) Measurement of RelA and RelB protein levels in *RELA RELB* double KO single cell clones generated in RPE1-hTERT cells.

**Figure S6.**
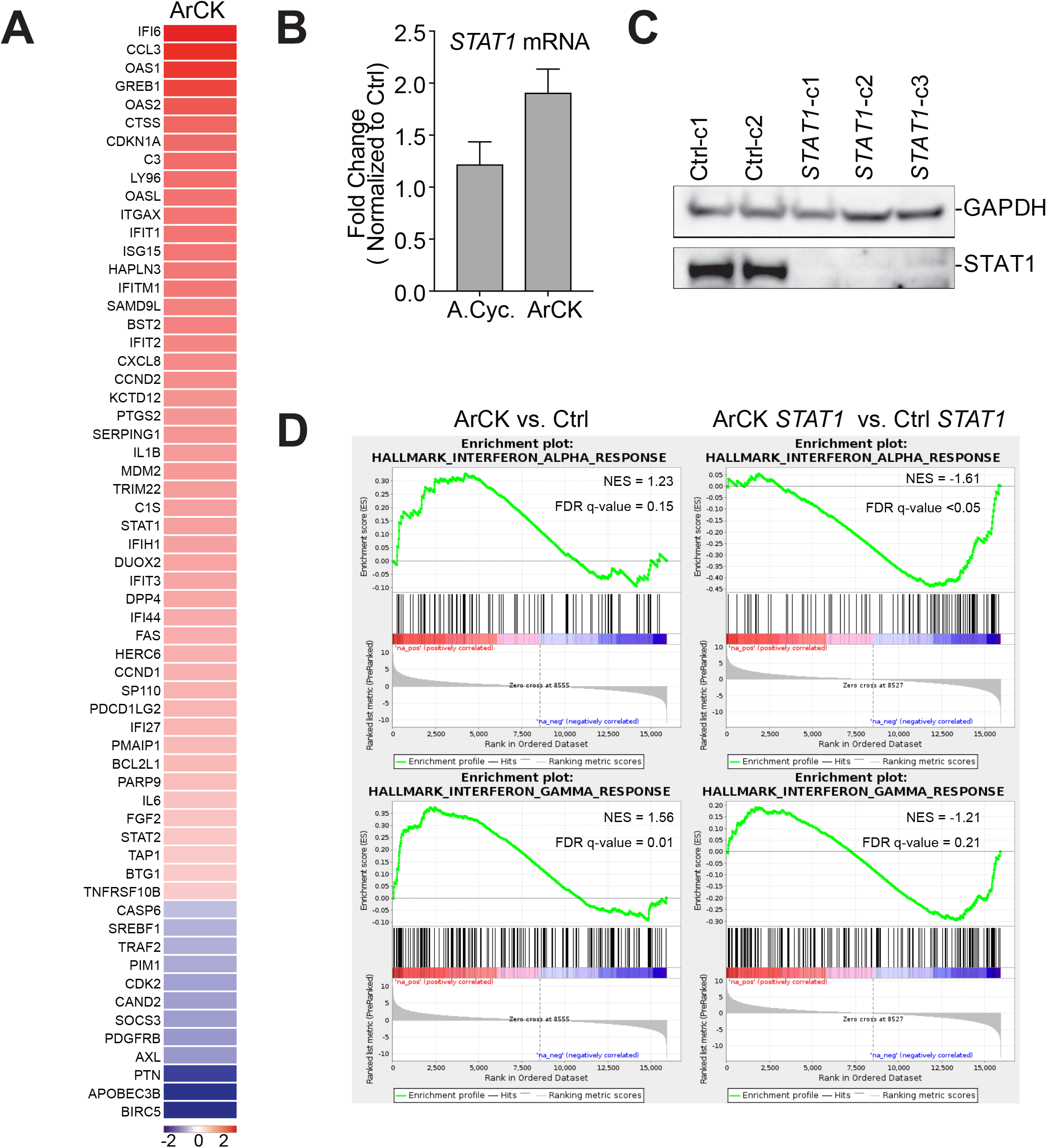
STAT1-responsive genes are induced in aneuploid cells. (A) Significantly differentially expressed STAT1 target genes in ArCK cells compared to euploid control cells (Log2 fold change, *p* value < 0.05). (B) Measurement of *STAT1* mRNA levels in aneuploid cycling (A.Cyc.) and ArCK cells compared to euploid control cells (n=3; mean ± SD). (C) Measurement of STAT1 protein levels in *STAT1* KO single cell clones generated in RPE1-hTERT cells. (D) GSEA enrichment plot for interferon alpha and interferon gamma hallmarks in ArCK cells generated in either control or *STAT1* RPE1-hTERT cells. Two single cell clones of control cell lines and three single cell cloned *STAT1* KO cell lines (shown in C) were used in the RNA-seq analysis.

**Figure S7.**
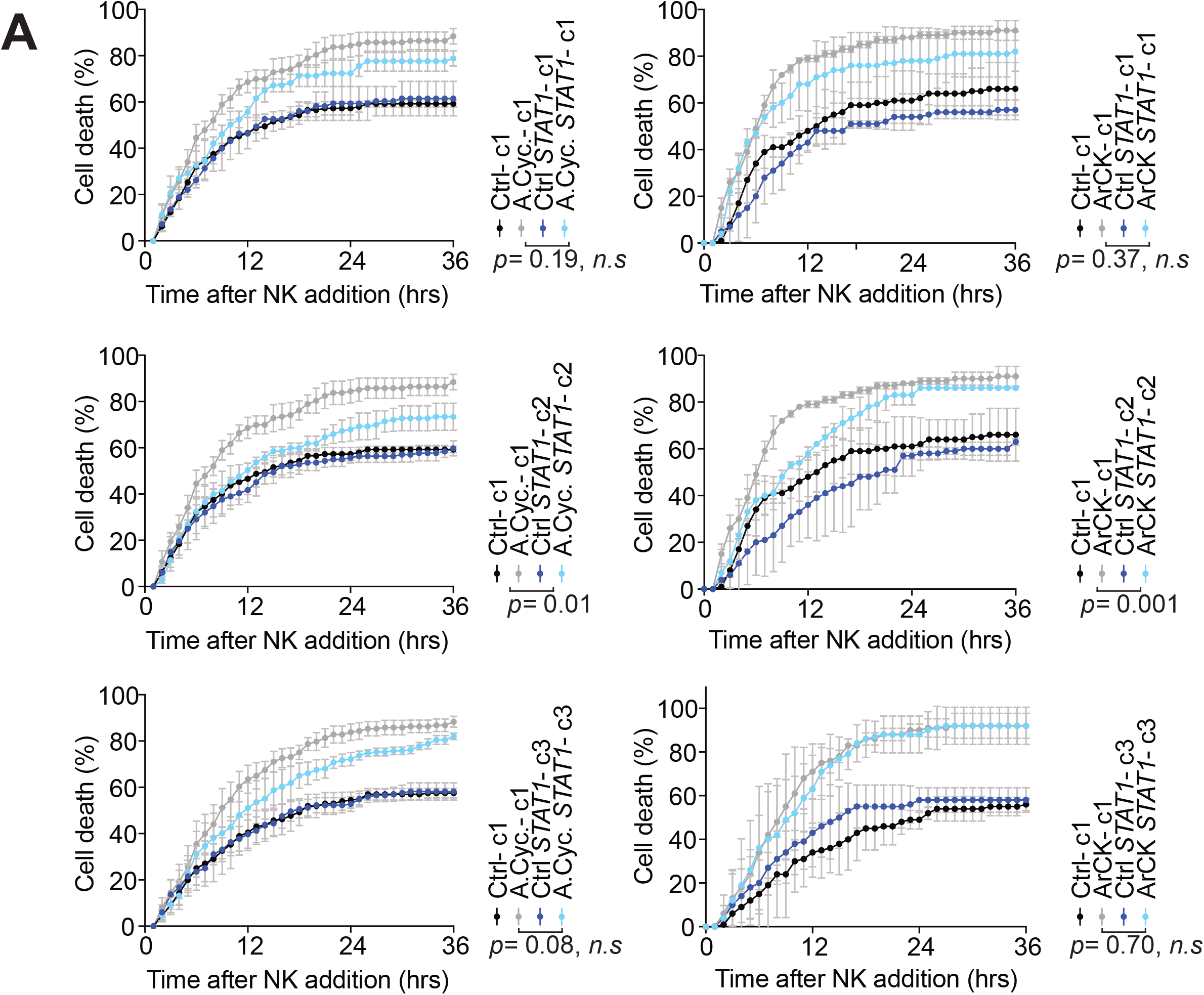
STAT1 inactivation does not affect NK cells-mediated killing of aneuploid cells. (A) *STAT1* KO RPE1-hTERT cells were generated using CRISPR-Cas9 method and single cell clones were obtained. Aneuploid cycling (A.Cyc.) and ArCK cells lacking *STAT1* were generated as described in Figure 1A and E. NK cell-mediated cell death was compared to control single cell clone which harbors an empty vector as described in Figure 1B (n=3; mean ± SD; *n.s*-not significant). The behavior of three different clones is shown in figure.

**Figure S8.**
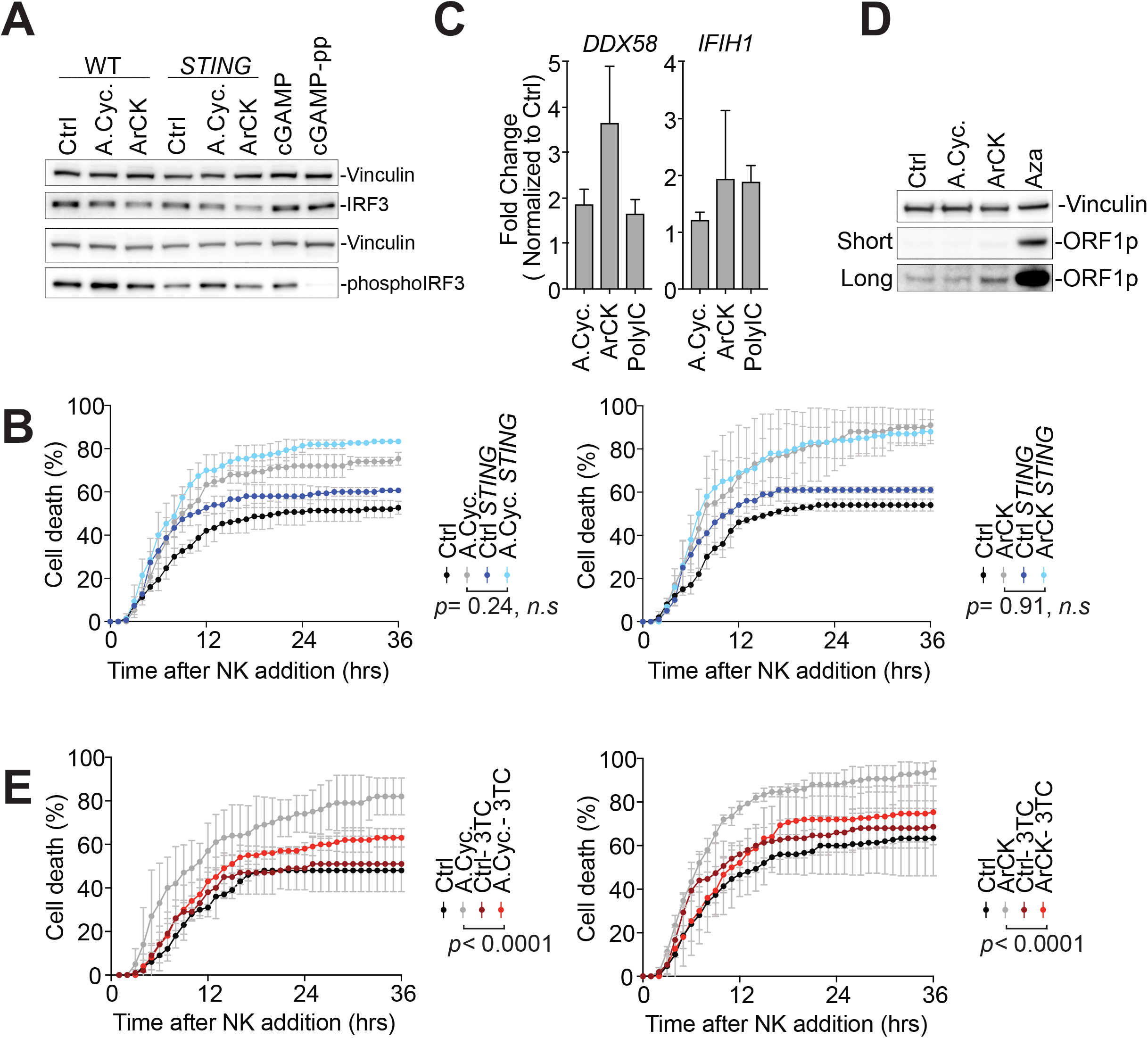
cGAS-STING and retrotransposon activity in aneuploid cells. (A) IRF3 and phospho-IRF3 levels were analyzed by western blot in euploid proliferating (Ctrl), aneuploid cycling (A. Cyc.), or ArCK cells that are either functional for STING (lanes 1-3) or lacking the gene (lanes 4-6). RPE1-hTERT cells treated with cGAMP (10 μg/ml) for 24hrs were used as a positive control (lanes 7 and 8). To confirm the specificity of the phospho IRF3-antibody, protein lysates from cGAMP treated cells were incubated with lambda phosphatase (lane 8). (B) *STING* KO RPE1-hTERT cells were generated using CRISPR-Cas9 method. Aneuploid cycling (A.Cyc.) and ArCK *STING* KO cells were generated and NK cell-mediated cytotoxicity was compared to control cell lines that harbor an empty vector as described in Figure 1B (n=2; mean ± SD). (C) Measurement of *DDX58* and *IFIH1* mRNA levels in aneuploid cycling (A. Cyc.) and ArCK cells by RT-qPCR shown as fold change compared to euploid control cells. RPE1-hTERT cells treated with PolyIC (10 μg/mL) for 24hrs were used as a positive control (n=3; mean ± SD). (D) ArCK and Aneuploid cycling (A.Cyc.) cells were grown as described in Figure 1A and E to determine ORF1 protein levels. RPE1-hTERT cells treated with azacitidine (5 μM) for 5 days (lane 4) was used as a positive control (n=2). (E) The effect of inhibiting reverse transcriptase activity on NK cell-mediated cytotoxicity in aneuploid cells. Aneuploid cycling (A.Cyc.) and ArCK cells were generated as described in Figure 1A and E. Aneuploid cycling and ArCK cells were continuously treated with reverse transcriptase inhibitor 3TC (7.5 μM) following chromosome mis-segregation. Control RPE1-hTERT cells were treated with 3TC for 3 days (timeline corresponds to aneuploid cycling cells treatment scheme). 3TC was washed out during the NK cell co-culture assay (n=3; mean± SD). Statistical analyses were performed as in Figure 1B.

## STAR METHODS

### RESOURCE AVAILABILITY

#### Lead Contact

Further information and requests for resources and reagents should be directed to and will be fulfilled by the Lead Contact, Stefano Santaguida.

#### Materials Availability

The knock out cell lines generated in RPE1-hTERT cells are available upon request.

#### Data and software availability

The RNA-seq data sets generated for this study can be accessed at Gene Expression Omnibus (GEO) database. The single cell DNA sequencing data can be accessed at Sequence Read Archive (SRA). The access numbers will be available upon publication.

### EXPERIMENTAL MODEL AND SUBJECT DETAILS

RPE1-hTERT cells were obtained from ATCC (ATCC Cat# CRL-4000) and HeLa cells (ATCC Cat# CCL-2). They were both cultured in Dulbecco’s modified Eagle’s medium (DMEM, Invitrogen) supplied with 10% FBS (Atlanta Biologicals of South America origin) and penicillin/streptomycin (100 U/ml) and L-Glutamine (2 mM).

Human primary IMR90 cells were obtained from ATCC (ATCC Cat# CCL-186). Adult and neonatal normal human dermal fibroblasts (NHDF-Ad and NHDF-Neo) were obtained from Lonza (Cat# CC-2511 and Cat# CC-2509, respectively). They were all cultured in Eagle’s Minimum Essential Medium (EMEM, ATCC) supplied with 10% FBS (Atlanta Biologicals) and penicillin/streptomycin (100 U/ml) and L-Glutamine (2 mM).

NK92-MI cells were obtained from ATCC (ATCC Cat# CRL-2408) and were cultured in MyeloCult H5100 medium (STEMCELL Technologies). All cells were grown at 37 °C with 5% CO2 in a humidified environment.

### METHOD DETAILS

#### Generation of aneuploid cycling cells and G1 arrested cells with complex karyotypes (ArCK)

To generate aneuploid cycling cells, 5 x 10^5^ RPE1-hTERT cells were plated on a 10cm culture dish and synchronized with thymidine (5mM, MilliporeSigma) for 24 hours. Cells were then released into complete medium. 6 hours after thymidine release, cells were switched into medium containing reversine (500nM, Cayman Chemical). Reversine was washed out after ~18 hours and aneuploid cycling cells were collected 48 hours after reversine washout.

To generate ArCK cells, 2.5 x 10^5^ RPE1-hTERT cells were plated on a 10cm culture dish and synchronized with thymidine (5mM) for 24 hours. Cells were then released into complete medium. 6 hours after thymidine release, cells were switched into medium containing reversine (500 nM). Reversine were washed out 18 hours later. 60 hours after drug washout, nocodazole (100 ng/ml, Sigma Aldrich) was added to the culture. 12 hours after nocodazole addition, mitotic cells were eliminated from the cell population by shake off. The shake-off process was performed for five times in total at a 12-hour interval. The cells left on the plate after five shake-offs were called the ArCK population.

#### Generation of cell-cycle arrested cells

To generate G1 arrested samples, 5 x 10^5^ RPE1-hTERT cells were plated on a 10cm culture dish and treated with the following drugs: doxorubicin (100 ng/ml, Sigma Aldrich), palbociclib (5 μM, LC Laboratories), and nutlin3 (10 μM, Cayman Chemical). Cells were collected after 7 days of drug treatment. For torin1 arrested samples, 1.5 x 10^6^ RPE1-hTERT cells were plated on a 10cm culture dish and were treated with torin1 (5 μM, LC Laboratories) for 24 hours.

#### Video Microscopy

All live cell imaging was performed using a spinning disk microscope (10x objective) with the environmental chamber maintained at 37 °C and 5% CO2 level. Target cells were plated onto 12-well glass bottom plates in complete normal growth medium at a density 4~6×10^4^ cells/well overnight to allow attachment. Target cells were switched into NK cell growth medium MyeloCult H5100 and incubated for 10 hours before starting the live cell imaging. To assess NK cell-mediated cell cytotoxicity, NK92-MI cells were re-suspended into target cell condition medium at the indicated NK cell to target cell ratio immediately before the start of the NK cell killing assay. The cell mixture was filmed for 36 hours at a 30min time interval. To assess target cell growth without NK cells, target cells were filmed using the same imaging setting and time scale.

#### Single cell sequencing

The experiment was performed as previously described (Knouse et al. 2014). Single cells were sorted into a 96-well plate and genomic DNA was amplified using the Single Cell Whole Genome Amplification Kit (Sigma). Amplified DNA was purified, barcoded, pooled, and sequenced on an Illumina HiSeq2000. Sequencing reads were aligned using BWA (0.6.1). HMMcopy (0.1.1) was used to estimate gene copy number in 500-kb bins. Cells with variability scores (VS) exceeding 0.34 were excluded from the analysis.

#### Cell cycle analysis by flow cytometry

Cells were trypsinized and resuspended into 10ml complete growth medium. After washing twice in cold phosphate-buffered saline (PBS), cells were fixed and permeablized in 70% ethanol at −20 °C overnight. Cells were then pelleted and incubated with ribonucelase A (2 mg/mL, Sigma Aldrich) and propidium iodide (100 μg/ml, ThermoFisher) for 1h at 37 °C immediately before flow cytometry analysis.

#### Cell volume analysis

Cultured cells were trypsinized from 10cm culture dish at ~70% confluency and were resuspended into normal complete growth medium immediately before measurement. 1ml of cell culture suspension were then added to isotonic buffer and the volume were measured on a Beckman coulter counter. At least 5000 cells were measured for each condition and the local mode for cell volume were presented in the graph.

#### Immunofluorescence

RPE1-hTERT cells were seeded onto fibronectin (10 μg/ml, Sigma Aldrich) coated coverslips at ~50% confluency and allowed to attached overnight. Prior to staining, cells were fixed at room temperature with either cold methanol (−20 °C) or 4% paraformaldehyde in PBS (4 °C) for 15mins. Cells were then permeabilized with 0.1% triton-X100 in PBS for 10mins and blocked in 4% bovine serum albumin (BSA) in PBS for 30mins. Primary antibodies were diluted in 4% BSA-PBS solution and incubated with cells for 90mins at room temperature. For fluorescence detection, Alexa Fluor 488 and 647 (Thermo Fisher) were used as secondary antibodies at a 1:1000 dilution in 4% BSA-PBS solution. Hoechst (100 ng/ml) was used to stain DNA. The following primary antibodies were used in experiments: anti-Phospho-histone H2A.X (Cell Signaling Technology #9718, 1:500), anti-Ki67 (Abcam 16667, 1:500).

#### Western Blot analysis

To prepare protein samples, protease inhibitor cocktail (Roche) and phosphatase inhibitor cocktail (Roche) were added to RIPA lysis buffer (Thermo Fisher Scientific) immediately before use. Cells were lysed in cold RIPA buffer and the lysate concentration was measured by either Bradford assay or Protein Assay Dye Reagent Concentrate (Bio-Rad, 5000006). The lysate was then diluted with loading buffer and heated at 98°C for 5 min. Proteins were resolved on either NuPAGE 4-12% Bis-Tris gels (Thermo Fisher Scientific) or 4–20% Criterion TGX Precast gels (Bio-Rad, 5678094) based on the manufacturer’s instructions and transferred onto 0.2 μm PVDF or nitrocellulose membranes. Blots were blocked for 30 mins at room temperature in TBS-Tween (0.1%) with either 5% BSA or 5% non-fat milk or OneBlock blocking buffer (Genesee Scientific). Primary antibodies were incubated over night at 4°C. The following primary antibodies were used: anti-GAPDH (Santa Cruz sc-365062 or sc-32233, 1:1000), anti-Vinculin (Sigma-Aldrich V9131, 1:5000), anti-Histone H3 (Cell Signaling Technology #4499, 1:1000), anti-p53 (Santa Cruz sc-126, 1:200), anti-p21 (Cell Signaling Technology #2947, 1:1000), anti-IκBα (Cell Signaling Technology #4814, 1:1000), anti-Phospho-IκBα Ser32/36 (Cell Signaling Technology #9246, 1:1000), anti-p65/RelA (Cell Signaling Technology #8242 or Santa Cruz sc-8008, 1:1000), anti-RelB (Abcam #180127, 1:1000), anti-Stat1 (Cell Signaling Technology #9175, 1:500), anti-IRF3 (Cell Signaling Technology, #11904, 1:1000), anti-Phospho-IRF3 (Cell Signaling Technology, #4947, 1:1000), anti-ORF1p (Cell Signaling Technology #88701, 1:1000).

#### Beta galactosidase staining

ArCK, aneuploid cycling, doxorubicin, palbociclib, nutlin3 and untreated euploid proliferating RPE1-hTERT cells were plated with normal complete growth medium into 6-well plates at a density of 4×10^5^ cells/well. Cells were left overnight to allow attachment to the plate. Samples were then fixed and stained for the β-Galactosidase activity using a senescence β-Galactosidase staining kit (Cell signaling technology #9860) following manufacturer’s instructions.

#### Cytokine measurement

ArCK, aneuploid cycling, doxorubicin, palbociclib, nutlin3, torin1 and untreated euploid proliferating RPE1-hTERT cells were generated as described above. 8ml of complete normal growth medium was placed onto cells and incubated for 36 hours. The conditioned medium was harvested and cell debris was eliminated by centrifugation. To determine the levels of secreted cytokines, condition medium was incubated with proteome profiler human XL cytokine array (R&D Systems, ARY022B) and cytokine levels were measured based on manufacturer’s instructions. The levels of interferon alpha and interferon beta in the condition medium were determined using IFN alpha and IFN beta human ELISA kit (Thermo Fisher Scientific, 411001 and 414101 respectively). The total cell number from each sample was measured using a cellometer (Nexcelom). All cytokines, IFN alpha, and IFN beta readings were normalized to cell number.

#### RNA sequencing and data analysis

Total RNA was purified using RNeasy Mini Kits (QIAGEN) and sequenced on Illumina HiSeq2000. The RNAseq data were aligned to a transcriptome derived from the human hg38 primary assembly and an ensembl version 89 annotation with STAR version 2.5.3a (Dobin et al. 2013). Gene expression was summarized using RSEM version 1.3.0 (B. Li & Dewey 2011) and samtools/1.3 (H. Li et al. 2009). An integer count table for differential expression analysis and log2 transcripts per million (TPM) with a plus 1 offset for data visualization was prepared with Tibco Spotfire Analyst (version 7.11.1). Differential expression analysis was done with DESeq2 (version 1.24.0, (Love et al. 2014)) running under R (version 3.6.0). Pre-ranked Gene Set Enrichment Analysis (versions 2-3.0_beta2 or 4.0.3, (Subramanian et al. 2005)) was run using the DESeq2 Wald statistic as a ranking metric and gene set collections from MsigDB (versions 6.2 or 7.0, (Liberzon et al. 2015)).

#### Generation of knock-out cell lines using the CRISPR-Cas9 system

Lentiviral CRISPR-Cas9 plasmid LentiCRISPR_v2-Puro (Brett Stringer’s Lab) cloned with guide RNA (gRNA) designed by Feng Zhang’s lab from the Broad Institute targeting exons of human *RELA* (AGCGCCCCTCGCACTTGTAG), *RELB* (TCGCCGCGTCGCCAGACCGC), and *STAT1* (ATTGATCATCCAGCTGTGAC) were purchased from GenScript. CRISPRv2 constructs along with packaging plasmids pMD2.G (Addgene 12259) and psPAX2 (Addgene 12260) were transfected into 293FT cells (Thermo Fisher, Cat# R70007) using TransIT-LT1 transfection reagent (Mirus). Virus was collected and the lentiviral titer was estimated by Lenti-X GoStix Plus (TaKaRa). RPE1-hTERT cells were plated at ~60% confluency and infected at a MOI of 1 for *RELA, RELB*, or *STAT1* single knockouts, and at a MOI of 2 for *RELA RELB* double knockout. Virus was washed out 20 hours post infection and the non-infected cells were selected against by puromycin treatment. LentiCRISPR_v2-Puro vector without gRNAs was integrated into RPE1-hTERT cells to generate the control cell line. 2 days after puromycin selection, single cells were sorted into 96 wells and expanded as individual clones. The successful knockout of *RELA, RELB* or *STAT1* was verified by western blot analysis. All experiments described above were performed in at least two single cell knock-out clones.

#### RT-qPCR analysis

Total RNA was purified using an RNeasy Mini Kit (QIAGEN). RNA concentration was determined using a nanodrop. 750ng of RNA was used for reverse transcription reactions using SuperScript III First-Strand Synthesis SuperMix (Invitrogen). *STAT1* mRNA level and *GAPDH* mRNA levels were quantified by qPCR using SYBR Premix Ex Taq (TaKaRa) on Roche Light Cycler. Primer sequences: *STAT1*-Forward GATG I I ICATTTGCCACCATCCGTTTTC. *STAT1*-Reverse GGCGTTTTCCAGAATTTTCCTTTCTTCC. *GAPDH*-Forward CCATGTTCGTCATGGGTGTGAACCATG. *GAPDH*-Reverse CCACAGCCTTGGCAGCGCCAGTAGAGG. *DDX58*_Forward CGGCACAGAAGTGTATATTGGATGCATTC. *DDX58*_Reverse GGAAGCACTTGCTACCTCTTGCTCTTC. IFIH1_Forward CTGGGACTAACAGCTTCACCTGGTGTTG. IFIH1_Reverse GCATCTGCAATGGCAAACTTCTTGCATG.

#### Nuclear/cytoplasm fractionation

Cells were lysed on ice for 10 min in NP-40 lysis buffer (10 mM HEPES pH 7.9, 1.5 mM MgCl2, 60 mM KCl, 1 mM DTT and 0.1% NP-40) supplemented with protease and phosphatase inhibitors. The lysates were centrifuged at 3000 rpm at 4°C for 5 min. The supernatant was collected as the cytoplasm fraction. The pellets, which represents nuclear fraction, were washed three times with wash buffer (10 mM HEPES pH 7.9, 1.5 mM MgCl2, 10 mM KCl and 1 mM DTT) and centrifuged at 2000 rpm at 4°C for 5 min. After the washes, the pellets were lysed with the SDS lysis buffer (one part of 5% SDS, 0.15M HCl pH 6.8, 30% Glycerol; and three parts of 25 mM TRIS HCl pH 8.3, 50 mM NaCl, 0.5% NP-40, 0.5% Sodium Deoxycholate, 0.1% SDS) supplemented with protease and phosphatase inhibitors and then sonicated. Protein concentration was measured using the DC Protein Assay (Bio-Rad, 5000116).

#### NF-κB Secreted Alkaline Phosphatase (SEAP) Reporter Assay

A NF-κB Secreted Alkaline Phosphatase Reporter Assay Kit (Novus biologicals, NBP2-25286) was used to measure the secretion of the SEAP protein under the control of the NF-κB promoter. To generate NF-κB secreted alkaline phosphatase reporter cell line, RPE1-hTERT cells were plated in a six-well plate in 2 mL DMEM, containing 10% FBS, L-Glutamine and nonessential amino acids. After 12 hours of incubation at 37°C and 5% CO2, cells were transfected with 1 μg pNF-κB /SEAP plasmid, using 6 μL Lipofectamine 3000 (Invitrogen, L3000008) per well. After 10-12 hours of incubation, the medium was replaced and reversine (500 nM) or DMSO were added for 60 hours. The supernatant was then collected to measure the levels of SEAP according to the manufacturer’s instructions. The SEAP assay standard curve used to calculate the sample SEAP concentration, was generated by loading a serial dilution of SEAP standard on the same plate. The absorbance was measured after 30 min of incubation with the PNPP substrate, using a GloMax Explorer Multimode Microplate Reader (Promega).

#### CCLE data analysis

Gene expression data for the CCLE lines were obtained from DepMap 19Q1 DepMap release; www.DepMap.org). A ssGSEA signature (Subramanian et al. 2005) score was calculated for the Hallmark_TNFA_signaling_via_NFKB gene set. Aneuploidy scores were obtained from Cohen-Sharir et al. (manuscript in revision). The association between the signature score and the aneuploidy groups was assessed by linear regression analysis, using the R package limma (Ritchie et al. 2015). Significance was calculated by empirical-Bayes moderated t-statistics.

### QUANTIFICATION AND STATISTICAL ANALYSIS

Statistical analysis was performed using GraphPad Prism or R software. Details of the statistical tests were stated in the associated figure legends. Error bars represent SD.

### KEY RESOURCES TABLE

| REAGENT or RESOURCE | SOURCE | IDENTIFIER |
| --- | --- | --- |
| **Antibodies** |  |  |
| Anti-p53 | Santa Cruz Biotechnology | Cat# sc-126, RRID:AB_628082 |
| Anti-p21 | Cell Signaling Technology | Cat# 2947, RRID:AB_823586 |
| Anti-Vinculin | Sigma Aldrich | Cat# V9131, RRID:AB_477629 |
| Anti-Histone H3 | Cell Signaling Technology | Cat # 4499, RRID:AB_10544537 |
| Anti- Phospho-histone H2A.X | Cell Signaling Technology | Cat# 9718, RRID:AB_2118009 |
| Anti-Ki67 | Abcam | Cat# 16667, RRID:AB_302459 |
| Anti-GAPDH | Santa Cruz Biotechnology | Cat# sc-365062, RRID:AB_10847862 |
| Anti-GAPDH | Santa Cruz Biotechnology | Cat# sc-32233, RRID:AB_627679 |
| Anti-IkBα | Cell Signaling Technology | Cat# 4814, RRID:AB_390781 |
| Anti-Phospho-IkBα Ser32/36 | Cell Signaling Technology | Cat# 9246, RRID:AB_2267145 |
| Anti-p65/RelA | Cell Signaling Technology | Cat#8242, RRID:AB_10859369 |
| Anti-p65/RelA | Santa Cruz Biotechnology | Cat# sc-8008, RRID:AB_628017 |
| Anti-RelB | Abcam | Cat# 180127 |
| Anti-Stat1 | Cell Signaling Technology | Cat# 9175, RRID:AB_2197984 |
| Anti-IRF3 | Cell Signaling Technology | Cat# 11904, RRID:AB_2722521 |
| Anti-Phospho-IRF3 | Cell Signaling Technology | Cat# 4947, RRID:AB_823547 |
| Anti-ORF1p | Cell Signaling Technology | Cat# 88701, RRID:AB_2800129 |
| **Chemicals, Peptides, and Recombinant Proteins** |  |  |
| Fibronectin | Sigma Aldrich | F1141 |
| Thymidine | MilliporeSigma | C946T48 |
| Reversine | Cayman Chemical | 10004412 |
| Nocodazole | Sigma Aldrich | M1404 |
| Doxorubicin | Sigma Aldrich | D2975000 |
| Palbociclib | LC Laboratories | p-7744 |
| Torin1 | LC Laboratories | t-7887 |
| Nutlin-3 | Cayman Chemical | 10004372 |
| BMS-345541 | Sigma Aldrich | B9935 |
| TNF-α | Sigma Aldrich | T0157 |
| cGAMP | Sigma Aldrich | SML1232 |
| LPS | Sigma Aldrich | L2630 |
| 5-Aza-2'-deoxycytidine | Sigma Aldrich | A3656 |
| Poly(I:C) | Invitrogen | tlrl-pic |
| 2',3'-Dideoxy-3'-thiacytidine | Sigma Aldrich | L1295 |
| **Critical Commercial Assays** |  |  |
| Click-iT EdU Alexa Fluor 488 Imaging Kit | Invitrogen | C10337 |
| GenomePlex Single Cell Whole Genome Amplification Kit | Sigma Aldrich | WGA4-500RXN |
| Senescence β-Galactosidase Staining Kit | Cell Signaling Technology | 9860 |
| Proteome Profiler Human XL Cytokine Array Kit | R&D systems | ARY022B |
| IFN beta Human ELISA Kit | Thermo Fisher Scientific | 414101 |
| Interferon alpha (Cell culture) Human ELISA Kit | Thermo Fisher Scientific | 411001 |
| NF-κB Secreted Alkaline Phosphatase Reporter Assay Kit | Novus biologicals, | NBP2-25286 |
| **Deposited Data** |  |  |
| RNA-Seq | This paper | Available upon publication |
| Single Cell DNA-Seq | This paper | Available upon publication |
| **Experimental Models: Cell Lines** |  |  |
| RPE1-hTERT | ATCC | ATCC Cat# CCRL-4000, RRID:CVCL_4388 |
| HeLa | ATCC | ATCC Cat# CCL2, RRID: CVCL_0030 |
| NHDF-Ad | Lonza | Cat# CC-2511 |
| NHDF-Neo | Lonza | Cat# CC-2509 |
| IMR90 | ATCC | ATCC Cat# CCL-186, RRID:CVCL_0347 |
| NK92-MI | ATCC | ATCC Cat# CRL-2408, RRID:CVCL_3755 |
| 293-FT | Thermo Fisher Scientific | R70007 |
| **Oligonucleotides** |  |  |
| *STAT1*-Forward | This paper | N/A |
| *STAT1*-Reverse | This paper | N/A |
| *GAPDH*-Forward | This paper | N/A |
| *GAPDH*-Reverse | This paper | N/A |
| *DDX58*_Forward | This paper | N/A |
| *DDX58*_Reverse | This paper | N/A |
| *IFIH1*_Forward | This paper | N/A |
| *IFIH1*_Reverse | This paper | N/A |
| **Recombinant DNA** |  |  |
| LentiCRISPR_v2-Puro | Addgene | Cat# 52961, RRID:Addgene_52961 |
| LentiCRISPR_v2-Puro-*RELA* | GenScript | Cat# SC1805 |
| LentiCRISPR_v2-Puro-*RELB* | GenScript | Cat# SC1678 |
| LentiCRISPR_v2-Puro-*STAT1* | GenScript | Cat# SC1805 |
| pMD2.G | Addgene | Cat# 12259, RRID:Addgene_12259 |
| psPAX2 | Addgene | Cat# 12260, RRID:Addgene_12260 |
| Software and Algorithms |  |  |
| GraphPad Prism | N/A | RRID:SCR_002798 |
| MATLAB | N/A | RRID:SCR_001622 |
| R Project for Statistical Computing | N/A | RRID:SCR_001905 |

